# Diversity of developing peripheral glia revealed by single cell RNA sequencing

**DOI:** 10.1101/2020.12.04.411975

**Authors:** OE Tasdemir-Yilmaz, NR Druckenbrod, OO Olukoya, AR Yung, I Bastille, MF Pazyra-Murphy, A Sitko, EB Hale, S Vigneau, AA Gimelbrant, P Kharchenko, LV Goodrich, RA Segal

## Abstract

The peripheral nervous system responds to a wide variety of sensory stimuli, a process that requires great neuronal diversity. These diverse peripheral sensory neurons are closely associated with glial cells that originate from the neural crest (NC). However, the molecular nature and origins of diversity among peripheral glia is not understood. Here we used single cell RNA sequencing to profile and compare developing and mature glia from somatosensory lumbar dorsal root ganglia (DRG) and auditory spiral ganglia (SG). We found that the glial precursors (GPs) differ in their transcriptional profile and prevalence in these two systems. Despite their unique features, somatosensory and auditory GPs undergo convergent differentiation to generate myelinating and non-myelinating Schwann cells that are molecularly uniform. By contrast, although satellite glia surround the neuronal cell bodies in both ganglia, we found that those in the SG express multiple myelination-associated genes, while DRG satellite cells express components that suppress myelination. Lastly, we identified a set of glial signature genes that are also expressed by placode-derived supporting cells, providing new insights into commonalities among glia across the nervous system. This comprehensive survey of gene expression in peripheral glia constitutes a valuable resource for understanding how glia acquire specialized functions and how their roles differ across sensory modalities.

## Introduction

The peripheral nervous system (PNS) is comprised of sensory and autonomic ganglia that relay multiple types of sensory information to the central nervous system (CNS). Although molecular diversity in the neurons of these ganglia is well characterized (Cane and Anderson, 2009; Chiu et al., 2014; Hockley et al., 2019; Nguyen et al., 2017; Shrestha et al., 2018; Usoskin et al., 2015), it is unclear whether there is also molecular diversity among the glial cells that associate with these neurons. Like glia in the CNS, peripheral glia are closely associated with sensory neurons and influence their functions in a variety of ways. The major glial subtypes in the PNS are Schwann cells and satellite glia. Schwann cells surround the axons and have important roles in fast propagation of action potentials, metabolic and trophic support to neurons, debris clearance and axon regeneration after injury (Beirowski et al., 2014; Jessen and Mirsky, 2019; Monk et al., 2015; Nave, 2010; Nguyen et al., 2009). Myelinating and nonmyelinating Schwann cells, the two major subtypes of Schwann cells, wrap around axons forming myelin and Remak bundles, respectively (Jessen et al., 2015). Satellite glia envelop the sensory neuron cell bodies (Hanani, 2005; Pannese, 1981), providing metabolic and trophic support to the neurons, buffering extracellular ion and neurotransmitters, clearing cell debris and enabling neuromodulation (Berger and Hediger, 2000; Hanani and Spray, 2020; Huang et al., 2013; Tang et al., 2010; Vit et al., 2008; Wu et al., 2009). In addition, activation of satellite glia after peripheral tissue injury or inflammation contributes to pathological pain (Takeda et al., 2009), making this glial type an important target for therapeutics. In spite of these multiple critical functions carried out by glia in the normal and diseased PNS, the nature and extent of molecular variation among glial subtypes across the various peripheral sensory systems is largely unknown.

Across the PNS, different types of sensory neurons experience unique functional demands that suggest there are also important differences among the associated glia. For example, in the auditory system, neurons in the spiral ganglion (SG) communicate signals from the ear to the brain with high speed and precision. Accordingly, SG neuron cell bodies and peripheral axons are heavily myelinated by satellite glia and Schwann cells. By contrast, in the pseudo-unipolar neurons of the dorsal root ganglion (DRG) of the somatosensory system, signals bypass the cell body, which is instead surrounded by non-myelinating satellite glia. Further, the degree of neuronal diversity in each system varies. DRG neurons convey several distinct types of tactile sensory information to the CNS, including temperature, touch, pain and spatial position, and this is reflected in tremendous molecular and anatomical diversity of these neurons. SG neurons, on the other hand, encode one modality, sound, and fall into only four distinct subtypes. It is not yet known how the fundamental differences in the function and composition of neurons in these two systems might be influenced by, and in turn influence, distinctive glial specializations.

All of the Schwann cells and satellite glia in the PNS develop from a common embryonic source, neural crest cells (NCCs). While NCCs show regional specializations along the rostro-caudal axis (Soldatov et al., 2019), their glial progeny have not been examined for similar rostro-caudal differences. In the trunk, NCCs that migrate ventrally give rise to all glial subtypes in two waves starting at E11 (Jacob et al., 2014; Maro et al., 2004). Glial precursors (GP)s are derived from NCCs, associate with the nascent DRG neurons, and generate both satellite glia and immature Schwann cells, from which non-myelinating (NMSCs) and myelinating Schwann cells (MSCs) arise (Kastriti and Adameyko, 2017). During cochlear development, NCCs initially form a sheath that surrounds the SG and the central axons and intercalate closely with the SG peripheral processes as they extend to form radial bundles (RB). As the peripheral axons extend from the SG, between E11.5 and E12.5 in mouse, migratory NCCs and GPs associate with the growing peripheral axons (Sandell et al., 2014), and this continues as the peripheral axons grow through the mesenchyme to reach the hair cells between E14 and E15 (Druckenbrod et al., 2020). Notably, while NCC-derived GPs in the trunk develop in close association with NCC-derived DRG neurons, the NCC-derived cochlear GPs interact with SG neurons and hair cells that are produced from the otic placode (Fritzsch et al., 2015; Pavan and Raible, 2012; Whitfield, 2015).

In addition to Schwann cells and satellite glia, the cochlea also houses a specialized population of glia-like supporting cells. Much like the glia that associate with ciliated sensory cells in *C. elegans* (Shaham, 2015), supporting cells interdigitate with and support the sensory hair cells within the organ of Corti, which is the sensory epithelium for hearing (Appler and Goodrich, 2011; Kelly and Chen, 2009).

Moreover, supporting cells briefly retain stem cell-like properties postnatally and can produce hair cells, making them an important target for efforts to promote hair cell regeneration to treat congenital deafness (Groves, 2010; Stone and Cotanche, 2007). Supporting cells express some familiar glial markers, such as Plp1 (Gomez-Casati et al., 2010) and Sox10 (Watanabe et al., 2000). However, the extent to which these cells exhibit a glial-identity is not yet clear. Further, unlike the NCC-derived Schwann cells and satellite glia, supporting cells originate in the otic placode. Therefore, comparison of supporting cells with other peripheral glia can help define genes that are characteristic of the glial identity, uncover shared developmental mechanisms that promote this identity, and reveal how differences both in the embryonic source and local environment affect glial precursors and their glial progeny.

To better understand how glial development and function is shaped by interactions with diverse neurons and environments, we used single cell sequencing to characterize developing peripheral glia and supporting cells from the SG and organ of Corti of the auditory system and from lumbar DRG and sciatic nerve (SN) of the somatosensory system. We found that GPs from somatosensory and auditory ganglia differ from one another in their transcriptional profile. As development progresses, some mature glial subtypes retain region-specific signatures whereas other GPs converge on a common path to produce Schwann cells with few molecular differences. Moreover, while GPs persist in postnatal DRG they largely disappear from the mature SG, suggesting differing programs of adult glial renewal in these two sensory ganglia. Among the more mature glial types, both DRG and SG satellite glia share many characteristics with CNS astrocytes. However, these two types of satellite glial cells are quite distinct from one another in their gene signatures, demonstrating that satellite glia in the two ganglia have different functions. In contrast, DRG and cochlear Schwann cells fall into only two groups, the myelinating and non-myelinating Schwann cells, and each category of Schwann cell is remarkably uniform across these differing ganglia. Together these data provide a novel resource for understanding the development and functions of peripheral glia and the contributions of these distinct glial cells to the wide scope of sensory perception.

## Results

### Characterization of peripheral glial cell clusters using Conos

To profile peripheral glia diversity and the developmental progression as these cells emerge from the NC and otic placode, we performed single cell analysis of glial cells from two distinct regions, cochlea and lumbar DRGs+sciatic nerve (SN). Glia were defined by expression of Plp1, a proteolipid protein, which is expressed in all peripheral myelinating and non-myelinating glial cells and in supporting cells (Buchstaller et al., 2004; Gomez-Casati et al., 2010). In *PLP-EGFP* mice, peripheral glia in both ganglia as well as the supporting cells of the organ of Corti are labeled as early as E14, and this expression is maintained in adults (Figure 1A, Figure S1A, B). Plp1+ glial populations were isolated from *PLP-EGFP* mice at E14, E18, and P14, stages extending from the time that cells derived from NCCs have coalesced to form ganglia (E14) to early differentiation (E18) and acquisition of mature differentiation state (P14), when satellite glia and Schwann cells are closely associated with neurons and their axons. We manually dissected lumbar DRGs and the attached SN as well as the cochlea, which includes the SG, peripheral processes, and the cochlear duct where PLP-EGFP glial supporting cells reside. Tissues were collected from the same animals in parallel (104 animals in total, see methods for details of genotypes), followed by enzymatic dissociation (Figure 1B). After FACS purification of GFP+ cells (Figure S2), we used the Indrop platform to prepare single-cell RNA-seq libraries. This method encapsulates single cells into droplets in which reverse transcription is performed in a way that incorporates unique barcodes into the cDNA of each individual cell (Klein et al., 2015) (Figure 1B). As an internal control, we included a small number of FACS-purified neurons from the same tissues (Figure S2), as marked by expression of the *Ai14* tdTomato reporter driven by *Neurog1*^*cre*^ or *Neurog1*^*creERT2*^ (Koundakjian et al., 2007; Madisen et al., 2010; Quinones et al., 2010). A total of 52,323 cells were collected and analyzed (see methods for quality control metrics), encompassing glia from two tissues and at three developmental stages.

**Figure 1:**
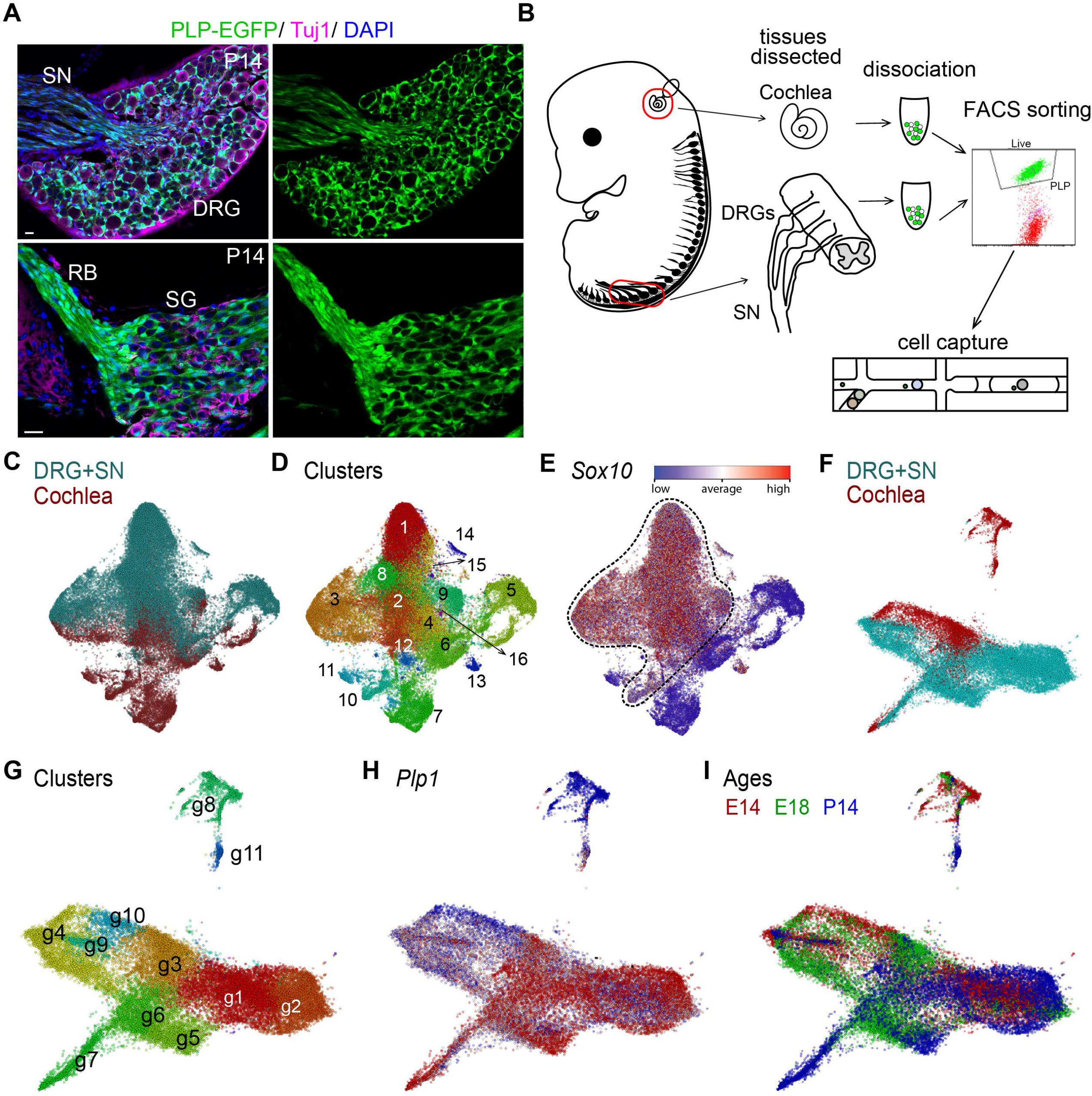
Single cell RNA-sequencing of peripheral glia from DRG+SN and cochlea at E14, E18 and P14. (A) P14 tissue sections of *PLP-EGFP* mice show labeling of DRG+SN and SG+RB satellite glia and Schwann cells. Tuj1 labels neurons, DAPI labels nuclei. Scale bars: 20um (B) Schematic of experimental design: Cochlea including SG, RB, and organ of Corti and lumbar DRGs+SN were dissected, dissociated, then cells were FACS sorted and loaded to InDrop chips for single cell collection. (C-E) Joint embedding (UMAP) of measured cells based on Conos dataset integration. Colors indicate (C) different tissue types (DRG+SN in blue, cochlea in red), (D) Individual clusters identified bioinformatically are labeled by numbers. (E) High Sox10 expression in all peripheral glia (highlighted by dotted line) (F-I) Joint embedding (largeVis) of glia after non-glia removed. Colors indicate (F) different tissue types (DRG+SN in blue, cochlea in red), (G) individual clusters labeled (g1-g11), (H) Plp1 expression in peripheral glia, (I) developmental stage (E14 in red, E18 in green, P14 in blue). DRG: dorsal root ganglion, SN: sciatic nerve, SG: spiral ganglion, RB: radial bundle.

To gain a comprehensive overview of the molecular relationships within this large population, we used the Conos algorithm (Barkas et al., 2019) to process the dataset collection. This approach allows us to overcome batch effects and establish correspondence between the cell subpopulations in different datasets. Conos constructs a global graph that compares gene expression of cells across different batches and from distinct locations and ages. In the combined graph including cells from distinct ganglia and developmental stages (Figure 1C), distance has meaning and cells with more similar gene expression profiles are closer to one another. We will refer to these combined graphs of measured cells based on Conos dataset integration as joint embeddings.

Using this approach, we identified 16 cell clusters (Figure 1D). As expected, two clusters contain the neuronal cells that were added, as identified by expression of the neuronal marker Tubb3 (cluster 5 and cluster 6), with SG neurons also labeled with Gata3 and DRG neurons also labeled with Ntrk1 (Figure S3). We also identified clusters corresponding to mesenchymal cells (clusters 7 and 11, Pou3f4), melanocytes (cluster 13, Mitf), macrophages (cluster 15, Fcgr3), and blood cells (cluster 16, Hba-a1) as identified by established markers (Figure 1D, Figure S3). These cells are likely contaminants from the FACS purification, with blood cells and macrophages from both ganglia and melanocytes and mesenchymal cells only from the cochlea (Figures 1C, 1D, Figure S3). Cochlear samples also include oligodendrocytes (Mobp+ cells in cluster 6, Figure S3), consistent with the fact that the dissected cochlea included a portion of the central nerve and its associated oligodendrocytes, which express a high level of Plp1 (Takeda et al., 2016) and were therefore expected to be captured by the FACS protocol. All other clusters (clusters 1-4, 8-10, 12, 14) are positive for Sox10, a peripheral glia marker (Kuhlbrodt et al., 1998) (Figure 1E). Of these, one cluster corresponds to glia-neuron doublets (cluster 14, Tubb3 and Sox10, Figure S3), predominantly derived from DRG. Thus, the Conos algorithm successfully groups cells according to known biological differences.

We computationally removed non-glial cell types, doublets, and oligodendrocytes and then designated the remaining clusters as peripheral glia (Figure 1E, peripheral glia shown with dashed line). This left a total of 29,307 cells, with 7,695 from the cochlea and 21,612 from lumbar DRGs+SN (Figure 1F). This difference in number presumably reflects the smaller number of glia in the cochlea compared to lumbar DRGs+SN. Analysis identified 11 clusters of peripheral glia (Figure 1G). All clusters show expression of the peripheral glia marker Plp1, though expression is lower in cluster 8 than the other clusters (Figure 1H). The majority of clusters included cells from both tissues and from multiple developmental stages (clusters g1, g3-7, g9, g10) (Figures 1F-I). Cluster identities were determined based on expression of known cell-cycle and glial genes. GPs are found in clusters g4, g9 and g10. GPs generate both satellite glia (clusters g1, g2, g11) and immature Schwann cells (clusters g3 and g6), from which mature non-myelinating Schwann cells (NMSCs) (cluster g5) and myelinating Schwann cells (MSCs) (cluster g7), arise. Cluster g8 represents supporting cells in the organ of Corti. These supporting cells are Plp1+ glia-like cells that originate from the otic placode and are clustered separately from the majority of NC-derived glia. Thus, this dataset includes peripheral glia from two developmental origins and captures their developmental trajectory from the progenitor stage to maturity in two sensory systems.

### Diversity of Glial precursors (GPs) from DRG and cochlea

GPs are descendants of the trunk and cranial NCCs; in turn GPs give rise to all mature glial subtypes. Unlike NCCs, GPs express genes characteristic of a glial lineage, such as Plp1. As expected for progenitor cells, we found that the most highly enriched genes in all GPs (clusters g4, g9 and g10, Figure 2A) include multiple cell cycle and proliferation-related genes. *Topoisomerase 2a* (*Top2a*), one of the top genes highly expressed in these clusters (Figure 2A), is important for chromosome condensation and segregation in dividing cells (Liu et al., 2019a; Milde-Langosch et al., 2013). Other genes associated with proliferation that are highly expressed in these clusters include *Cyclin Dependent Kinase 1* (*Cdk1*) and *cyclin A2* (*Ccna2*) (Figure S4). GP clusters also express genes with dual functions in proliferation and DNA repair/apoptosis, including *Birc5* (*Baculoviral IAP Repeat Containing 5*) and *Hmgb2* (*High Mobility Group Box 2*) (Li et al., 2018; Nagaki et al., 1998; Pusterla et al., 2009; Vequaud et al., 2016) (Figure S4). Other than the GPs, only a few cells in cluster g8 (supporting cells) express proliferation-related genes (Figures 1G, 2A, Figure S4); these g8 proliferative cells may represent placodal precursors that generate supporting cells.

**Figure 2.**
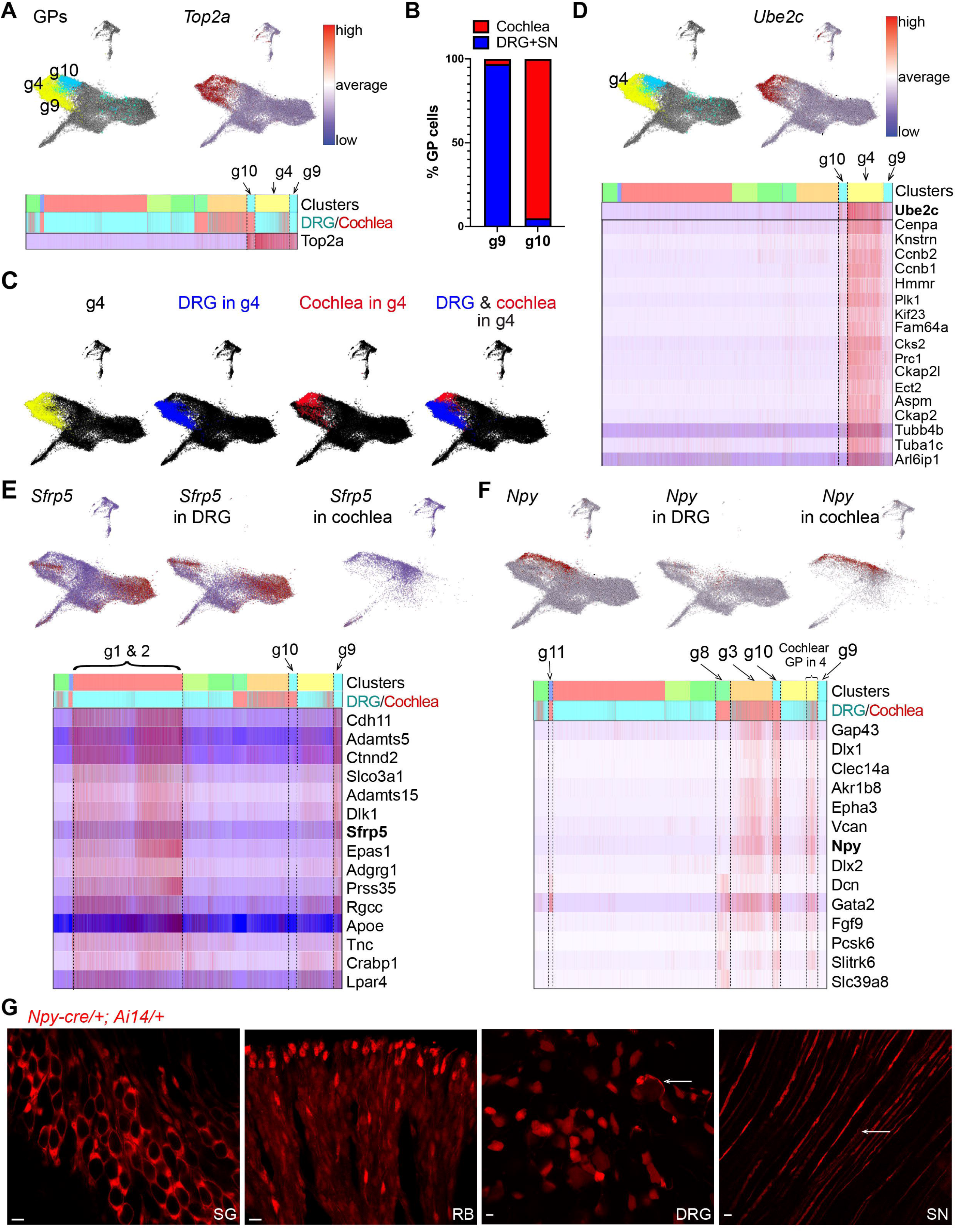
Glial precursors (GPs) from DRG and cochlea have distinct expression profile. (A) In Joint embedding (largeVis) of glia, GP clusters g4, g9 and g10 are labeled in yellow, cyan and blue, respectively. Joint embedding and heatmap show Top2a expression in all GP clusters. (B) Graph showing g9 contains %97 DRG+SN GPs and g10 contains %95 cochlear GPs. (C) Joint embeddings (largeVis) of glia: DRG GPs in blue, cochlear GPs in red. DRG and cochlear GPs do not intermix within cluster g4. (D) Joint embedding (largeVis) shows high Ube2c expression in cluster g4. Heatmaps show top genes expressed highly in cluster g4 compared to g9 & g10. (E) Sfrp5 is enriched in DRG+SN glia compared to cochlear glia. Joint embeddings (largeVis) show Sfrp5 expression in all glia, in DRG+SN glia and cochlear glia from left to right. Heatmaps show top genes expressed highly in cluster g9 compared to g10. Npy is enriched in cochlear glia compared to DRG+SN glia. Joint embeddings (largeVis) show Npy expression in all glia, in DRG+SN glia and cochlear glia from left to right. Heatmaps show top genes expressed highly in cluster g10 compared to g9. (G) Whole mount images from 2-month-old *Npy*^*Cre*^/+; *Ai14*/+ mice show *Npy*^*Cre*^ labels satellite glia in SG and Schwann cells in RB, but labels neurons in DRG+SN. Arrows indicate rare *Npy*^*Cre*^ labeling in DRG & SN glia. Scale bars: 10um.

Within the joint embeddings, GPs from distinct ganglia are separated from each other, suggesting that they express different genes depending on where they are located. Although all GP clusters express genes associated with proliferation, cluster g9 contains predominantly DRG+SN GPs with only 3% derived from the cochlea, while cluster g10 is predominantly cochlear GPs with only 5% from the DRG+SN (n= 1752 cells, 104 mice) (Figure 2B; Figures S5A, SB show the cells in joint embeddings). In addition, although cluster g4 includes cells from both tissues, cells segregate within the cluster according to tissue type (Figure 2C). Differential gene expression analysis across the three clusters revealed enhanced expression of proliferative signature genes in cluster g4 compared to clusters g9 and g10, including *Cdk1, Ccna2, Birc5*, and *Ube2C* (*Ubiquitin Conjugating Enzyme E2 C*), which encodes a ubiquitinating enzyme critical for regulated destruction of mitotic cyclins enabling cell cycle progression (Figure 2D, Figure S4). Other highly expressed genes in cluster g4 also have critical roles in the cell cycle, such as *Cenpa, Knstrn, Plk1* and *Kif23* (heatmap in Figure 2D) (Barr et al., 2004; Fang et al., 2009; Hutterer et al., 2009; Sullivan, 2001). This signature suggests that the shared GP cluster g4 represents an actively dividing, early pool of GPs that subsequently engender more specialized GPs that populate the DRG+SN (cluster g9) and cochlea (cluster g10).

To learn more about the molecular features that might distinguish GPs developing in the DRG *vs* cochlea, we compared gene expression between clusters g9 and g10, which will be referred to as DRG and cochlear GPs, respectively. *Sfrp5*, which encodes a Wnt inhibitor, is one of the top genes highly expressed in DRG GPs compared to cochlear GPs (Figure 2E). *Sfrp5* and other DRG GP-enriched genes are also highly expressed in DRG satellite glia (clusters g1 & g2). As almost all of these cells came from the DRG (Figure 2E), these data suggest that the selective gene set expressed in DRG GPs is characteristic of DRG GPs and many of their descendants. Similarly, many genes distinctive for cochlear GPs, such as *Gata2, Npy* and *Epha3*, are shared with other cochlear specific clusters and are not highly expressed in lumbar glial cells (Figure 2F). For instance, *Dcn*, one of the top markers for the cochlear GPs, is also expressed in supporting cells (cluster g8), whereas *Gata2* is also expressed in cochlear satellite glial cells (cluster g11) (Figures 1F, 1G, 2F heatmap). Notably, the cochlear GPs in cluster g4 and cochlear immature Schwann cells in cluster g3 also express *Gata2, Npy*, and *Epha3*. These data suggest that at early stages, GPs from the DRG and cochlea are already headed along distinct developmental paths.

Genetic fate mapping studies confirmed the unique contributions of Npy+ cochlear GPs to the auditory system. *Npy* is enriched in cochlear GPs in clusters g4 and g10 compared to DRG GPs and is also more highly expressed in the cochlear immature Schwann cells in cluster g3 compared to immature Schwann cells from the DRG+SN (Figure 2F). Moreover, at E18, Npy is expressed in cochlear but not DRG glia. To validate this difference and also follow the fates of *Npy*-expressing cells *in vivo*, we examined 2 month old *Npy*^*Cre*^; *Ai14* adult mice (2 mice; n=3 fields for DRGs, n=4 fields for SN from 6 DRGs and 2 SN per animal; n=4 fields for cochlea). In the cochlea of these animals, tdTomato broadly labels satellite glia in the SG and Schwann cells lining the RB, as predicted by expression of *Npy* in cochlear GPs (Figure 2G). In contrast, few Schwann cells and satellite glia are labeled in the somatosensory system (Figure 2G, arrows), matching the small Npy+ population of DRG GPs. Instead, a subset of DRG neuronal cell bodies in DRG and their axons in SN are labeled, consistent with known expression of Npy in somatosensory neurons (Figure 2G) (Mantyh et al., 1994). These data show that Npy distinguishes cochlear GPs from DRG GPs and confirms that cochlear GPs give rise to cochlear satellite glia and Schwann cells. Thus, *Npy*^*Cre*^ mice provide a good tool to study cochlear glia starting at early developmental times. Together with the scRNA-seq results, *Npy*^*Cre*^ fate-mapping indicates that DRG and cochlea GPs are molecularly distinct populations that generate glia in a tissue-specific manner.

In addition to these molecular differences, the GPs that populate the DRG and cochlea also differ in their long-term proliferative properties. At E14, GPs are present in both the DRG+SN and the cochlea (Figure 3A). By contrast, at P14, GPs are almost absent from the cochlea (Figure 3A). Comparison of GP number in both tissues further illustrates the persistence of a proliferative population in DRG+SN and not in the cochlea (n=3391 cells, 53 mice) (Figure 3B). In support of this hypothesis, RNAscope, semi-quantitative fluorescent RNA *in situ* hybridization analysis, showed strong expression of the proliferative marker *Top2a* in a subset of PLP-EGFP+ glia in the DRG+SN, but not in the cochlea of P14 *PLP-EGFP* mice (n=4 fields from SG and 3 fields from RB, from 2 mice; n=16 fields each for DRG & SN from 6 DRGs and 2 SN per animal, from 5 mice) (Figure 3C). Similar results were observed at P22 (2 mice; n=4 fields each for SG & RB; n=14 fields for DRG and n=10 fields for SN from 6 DRGs and 2 SN per animal) (data not shown). To independently assess glial proliferation in these ganglia, we injected EdU into *PLP-EGFP* mice at P21 and then counted the number of Plp1+ glial cells that incorporated EdU at P22, as well as those that expressed the proliferation marker Ki67 (Figure 3D). The average number of EdU + and Ki67+ glial cells per ganglion and per volume was dramatically lower in the cochlea (EdU+ glia 1.4±1.4 ⨯10^−6^ in SG, 0.1±0.1 ⨯10^−6^ in RB; Ki67+ glia 1.8±2.3 ⨯10^−6^ in SG, 0±0 ⨯10^−6^ in RB, each value um^-3^, mean ± SD) compared to lumbar DRG+SN (EdU+ glia 7.2±2.1 ⨯10^−6^ in DRG, 4.4±1.0 ⨯10^−6^ in SN; Ki67+ glia 13.7±2.6 ⨯10^−6^ in DRG, 5.5±1.4 ⨯10^−6^ in SN, each value um^-3^, mean ± SD) (6 mice; n=21 fields each for SG & RB from 8 ears for cochlea; n=30 fields each for DRG and & SN from 6 DRGs and 2 SN per animal) (Figures 3E, 3F). Thus, glia maintain the ability to proliferate in the DRG+SN but not in the cochlea, a difference that may have important implications for glial plasticity in these two regions.

**Figure 3:**
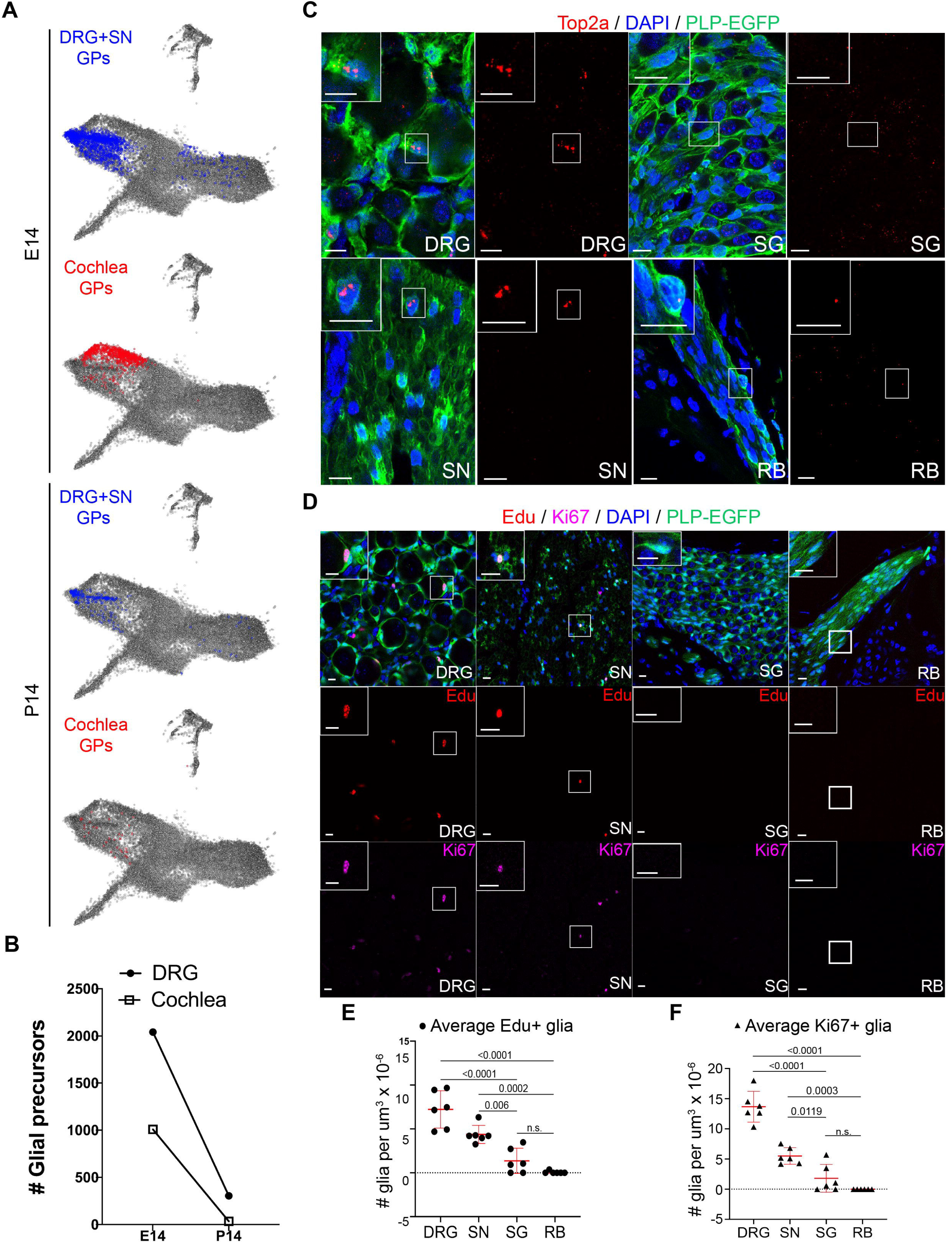
Glial precursors (GPs) are more prevalent and enduring in DRG than in cochlea. (A) Joint embeddings (largeVis) of glia, all cochlear GPs in red, all DRG GPs in blue. At P14, some GPs in DRG are still present, but almost none remain in cochlea. (B) Graph shows number of GPs from DRG and cochlea at E14 and P14 from (A). (C) RNAscope for Top2a on P14 tissue sections of *PLP-EGFP* mice. Strong Top2a puncta in glia from DRG+SN, but not from SG+RB. Insets are high mag views. (D) Edu and Ki67 stains in DRG, SN, SG and RB on tissue sections of *PLP-EGFP* mice. Edu injections at P21, tissue collected at P22. (E-F) Quantification of Edu (E) and Ki67 (F) stains from (C). Number of EdU+/Ki67+ and Plp1+ glia divided by tissue volume. p values indicate the results of Tukey’s test following one-way ANOVA. Error bars represent ± SD. Scale bars: 10um

### Characterization of Schwann cell subtype clusters

Schwann cells are one of the main glial subtypes that are derived from GPs throughout the PNS. Given the differences between DRG and cochlear GPs, we wondered whether there were also differences in Schwann cell development. Throughout the PNS, GPs give rise to immature Schwann cells, which differentiate either into NMSCs or promyelinating Schwann cells that subsequently develop into MSCs (Jessen and Mirsky, 2005). Promyelinating Schwann cells associate with axons in a 1:1 relationship but have not differentiated fully to generate a myelin sheath. To delineate the path that GPs in the DRG+SN and cochlea follow to produce these different Schwann cell types, we looked for markers associated with Schwann cell development and maturation.

We found two clusters of immature Schwann cells (cluster g3 and g6), as defined by expression of Ngfr and L1cam (Figure 4A) (Jessen and Mirsky, 2005). 90% of the cells in cluster g6 originated in the DRG+SN and thus are mostly DRG+SN immature Schwann cells (Figures 1F, 4A). Cluster g3 shows high expression of another immature Schwann cell marker Gap43 (Jessen and Mirsky, 2005) and largely consists of cochlear immature Schwann cells, with 73% cells in the cluster deriving from the cochlea (Figures 1F, 4A). Of all immature Schwann cells identified (cluster g3 & g6), 93% of those that came from the cochlea reside in cluster g3. Consistent with the identification of cluster g3 as a cochlear cell type, differential gene expression analysis showed that genes highly expressed in cluster g3 compared to the rest of the cells include the gene set characteristic of cochlear GPs, including *Gata2* and *Npy* (Figure S6). Thus, GPs in each ganglion produce immature Schwann cells that retain tissue-specific characteristics.

**Figure 4:**
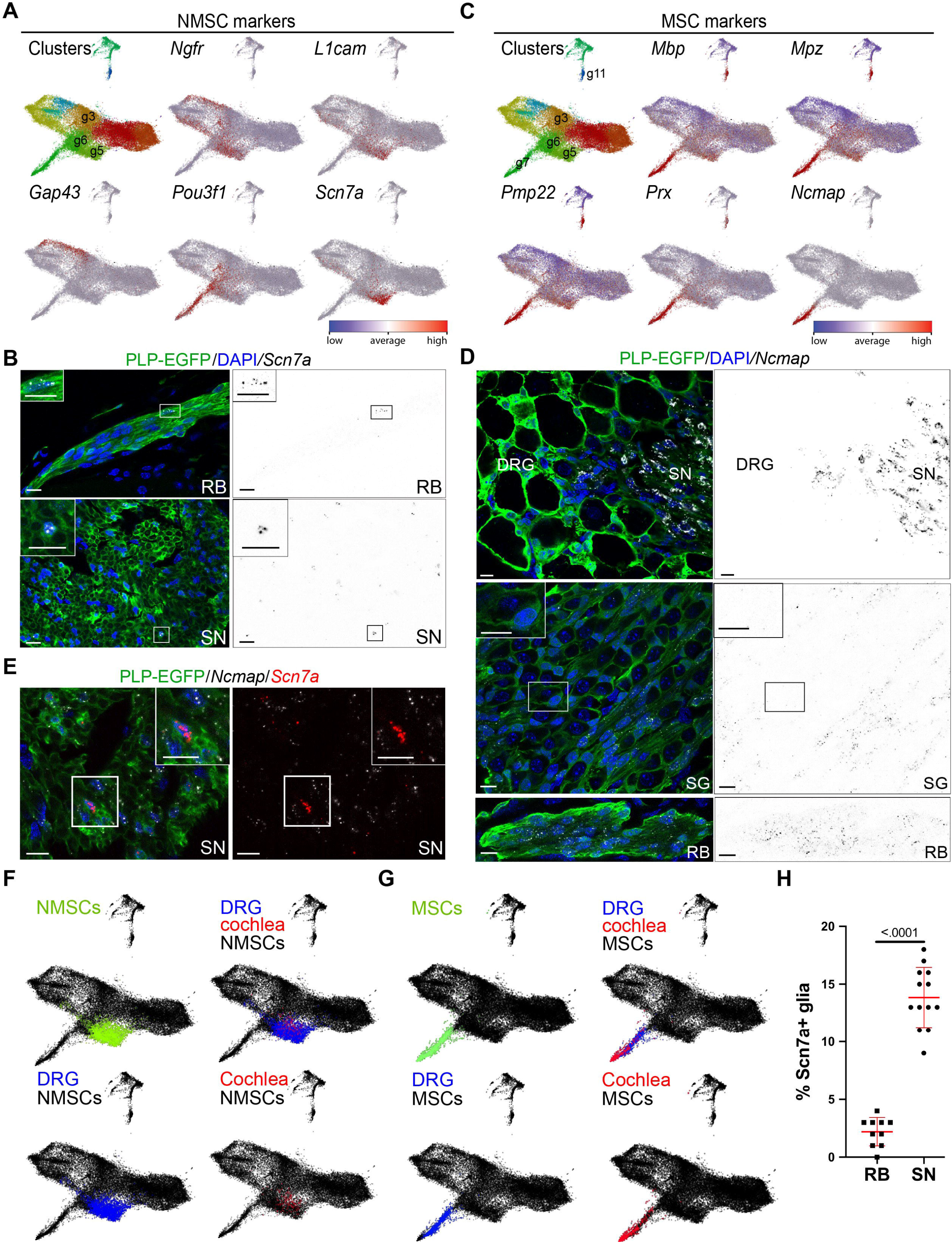
Schwann cell subtypes have consistent gene signatures irrespective of location. (A, C, F, G) Joint embeddings (largeVis) of glia show (A) expression of markers of immature Schwann cell (Ngfr, L1cam and Gap43), promyelinating Schwann cell (Pou3f1), NMSC (Ngfr and L1cam), and Scn7a, (C) expression of MSC markers Mbp, Prx, Mpz, Pmp22, and Ncmap. (A) Scn7a is most specific for NMSCs. (B-E) PLP-EGFP labels glia, DAPI labels nuclei. Scale bars: 10um (B) RNAscope on P14 sections show Scn7a puncta in Plp1+ Schwann cells in SN and RB. Scn7a is in white on the left, and black on the right panels (color inverted). (C) Ncmap is the most specific marker for MSCs. (D) RNAscope on P14 sections shows Ncmap puncta in Plp1+ Schwann cells in nerves next to DRG and SG, but not in satellite glia in DRG or SG. Ncmap is in white on the left, and black on the right panels (color inverted). Ncmap puncta seen in SG are localized in Schwann cells around the nerves. Inset shows absence of puncta in SG satellite glia. (E) RNAscope on P14 sections shows Scn7a and Ncmap puncta in non-overlapping Plp1+ Schwann cells. (F) DRG and cochlear NMSCs intermingle within NMSC cluster. DRG and cochlear MSCs intermingle within MSC cluster. (H) Graph showing % Scn7a+ glia is significantly higher in SN compared to RB. Average number of Scn7a+ glia/all cells is 2.2% in the RB whereas 13.85% in the SN (two-tailed unpaired t-test, p<.0001). Error bars represent ± SD.

Immature Schwann cells differentiate into MSCs or NMSCs that can be difficult to distinguish from their precursors due to the lack of cell type-specific markers. For instance, as well as marking immature Schwann cells, *Ngfr* and *L1cam* are also expressed by NMSCs (Jessen and Mirsky, 2005). Similarly, in our dataset, *Ngfr* and *L1cam* are expressed by cells in multiple clusters, including DRG and cochlear GPs and DRG sensory neurons, immature Schwann cells from each tissue (clusters g3 and g6), and cluster g5, which contains both DRG and cochlear cells (Figures 1G, 4A, Figure S7A). Cells in clusters g3 and g6 also express the promyelinating Schwann cell marker *Pou3f1* (Arroyo et al., 1998; Jessen and Mirsky, 2005; Zorick and Lemke, 1996) (Figure 4A). By contrast, the top gene enriched in cluster g5 is *Scn7a/Nav2*.*1*, which encodes an atypical Na-sensitive Na-channel known to be expressed by NMSCs and not MSCs (Hiyama et al., 2002; Watanabe et al., 2002) (Figure 4A). This suggests that cluster g5 corresponds to NMSCs, whereas clusters g3 and g6 contain immature Schwann cells able to generate both NMSCs and MSCs. Consistent with the fact that immature Schwann cells develop before NMSCs, more P14 cells are found in cluster g5 (823 cells) than in cluster g6 (521 cells) and cluster g3 (691 cells).

Furthermore, the level of *Scn7a* increases with time, suggesting that cells in cluster g5 that express lower *Scn7a* levels represent immature NMSCs (Figure S7B). *Scn7a* seems to be selective for NMSCs among the peripheral glia, although it is also expressed in some neurons, albeit at much lower levels than *Ngfr* and *L1cam* (Figure 4A, Figure S7A). Indeed, RNAscope confirmed expression of *Scn7a* in a subset of Schwann cells associated with the peripheral nerves (SN and RB), with no expression in satellite glia in either the DRG or SG (n=10 fields each for SG & RB from 4 ears for cochlea, 4 mice; n=13 fields each for DRG & SN from 6 DRGs and 2 SN per animal, 6 mice) (Figure 4B, Figure S7C). Thus, Scn7a reliably marks NMSCs in both sensory systems.

Immature Schwann cells also give rise to MSCs, which are defined primarily based on expression of genes required for myelin formation. However, our data showed that expression of myelin-related genes is not limited to one cluster. Instead, we observed broad expression of genes such as *Mbp, Mpz, Pmp22*, and *Prx* (Siems et al., 2020), with high levels in clusters g7 and g11 and low levels in multiple other clusters (Figure 4C). Cluster g7 is composed of both DRG and cochlear glia, whereas cluster g11 only contains cochlear glia (Figures 1F, 1G). Cluster g11, in addition to genes characteristic of myelinating cells, also expresses several markers for satellite glia, including *Glul* (Figure 5A) and *Kcnj10/Kir4*.*1* (Figure 5B). Since satellite glia myelinate cell bodies in the SG (Hibino et al., 1999), we defined cluster g11 as cochlear satellite glia and cluster g7 as MSCs.

**Figure 5:**
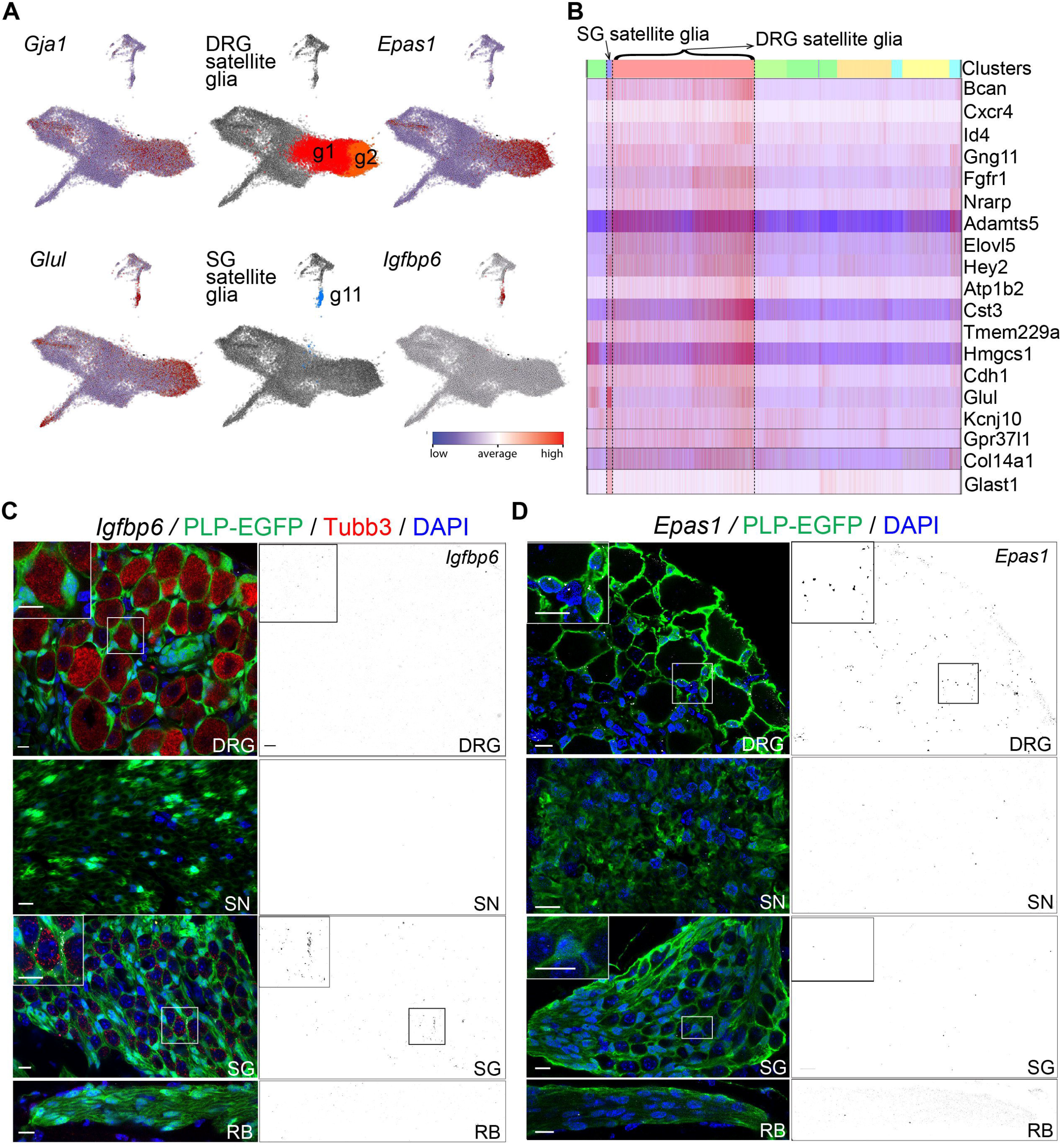
Satellite glia from DRG and cochlea cluster separately and express unique markers. (A) Joint embeddings (largeVis) show expression of satellite glia markers *Gja1* and *Glul*, clusters of DRG (g1&2) and SG (g11) satellite glia, new markers for DRG satellite glia (Epas1) and for SG satellite glia (Igfbp6). (B) Heatmaps of top genes shared between DRG and SG satellite glia. (C, D) RNAscope on P14 sections. PLP-EGFP labels glia, DAPI labels nuclei. Scale bars: 10um (C) Igfbp6 is expressed in satellite glia from SG, not from DRG. Igfbp6 puncta labeled in white on the left, black on the right panels (color inverted). (D) Epas1 is expressed in satellite glia from DRG, not from SG. Epas1 puncta labeled in white on the left, black on the right panels (color inverted).

Given the wide expression of genes associated with myelination, we sought to define more selective markers for MSCs. Differential analysis for genes enriched specifically and highly in cluster g7 compared to other peripheral glia identified *Ncmap* (*Non-Compact Myelin Associated Protein*) (Figure 4C).

RNAscope for *Ncmap* confirmed strong expression in Schwann cells and not in satellite glia in both ganglia (n=10 fields each for SG & RB from 4 ears for cochlea, 4 mice; n=19 DRG and 18 SN fields from 6 DRGs and 2 SN per animal, 11 mice) (Figure 4D). Furthermore, *Ncmap* expression does not overlap with expression of the NMSC marker *Scn7a*, consistent with the fact that these two genes are restricted to cluster g7 and g5, respectively (n=10 fields each for SG & RB from 4 for cochlea, 4 mice; n=13 fields each for DRG & SN from 6 DRGs and 2 SN per animal, 6 mice) (Figure 4E). Ncmap is a glycoprotein expressed preferentially in the myelin of PNS versus CNS and is localized mainly to the Schmidt-Lanterman incisures and paranodes of noncompact peripheral nerve myelin (Ryu et al., 2008). Since *Ncmap* expression is restricted to MSCs whereas other known MSC markers are expressed more generally, we suggest that *Ncmap* expression is likely to be a better marker for MSCs, and that this specificity can be exploited to identify and manipulate MSCs in a selective fashion.

Despite the fact that DRG and cochlear GPs are molecularly distinct, we found that NMSCs and MSCs from these two ganglia cluster together and are intermingled (Figures 4F, G). This suggests that even though Schwann cells arise from unique GPs and reside in different ganglia, their gene expression profile and function are remarkably uniform. The consistency among NMSCs and MSCs from distinct locations suggest that there is a singular set of genes that enables efficient axon wrapping and a second gene set that enables 1:1 myelinating insulation of axons. Thus, similar pathways seem to be active in Schwann cells from distinct tissues, overriding region-specific differences in GPs and immature SCs.

Although Schwann cells from the cochlea and SN are similar at the molecular level, the proportion of MSCs and NMSCs is quite different in each tissue. We observed in our dataset that the ratio of MSC:NMSC is 1:2.5 in the SN, whereas it is 4:1 in the cochlea. Thus, NMSCs account for 71% of the Schwann cells in the SN, but only 20% of the Schwann cells in the cochlea. Consistent with this difference in prevalence, RNAscope for *Scn7a* showed significantly fewer of the NMSCs associated with SGN peripheral processes in the RB (2.2±0.01%, value mean ± SD) compared to the SN (13.85±0.03%, value mean ± SD) (n=10 RB fields from 4 ears, 4 mice; n=13 SN fields from 2 SN per animal, 6 mice) (*P*<0.0001, two-tailed unpaired t-test, Figure 4H). The larger proportion of MSCs present in the cochlea is consistent with the need for high velocity transmission in the auditory system. The difference in the proportion of MSCs and NMSCs may reflect dissimilarities between the differentiation pathways from GPs to MSCs and NMSCs in the two ganglia, pathways that require Neuregulin-ErbB signaling between the axons and the immature Schwann cell precursors (Newbern and Birchmeier, 2010).

### Satellite glia in DRG and cochlea differ from one another

In contrast to the NMSCs and MSCs, satellite glia exhibit morphologic and functional differences depending on their location. Consistent with this diversity, three clusters expressed markers associated with satellite glia: g1, g2, and g11. As noted above, cluster g11 consists of SG satellite glia, based on expression of known markers, *Glul/GS* and *Kcnj10/Kir4*.*1* (Figures 5A, 5B). Cells in clusters g1 and g2 also expressed satellite glia markers *Gja1/Cx43* (*Connexin 43*) (Procacci et al., 2008), which was also observed in some GPs (Figure 5A). 97.5% of the cells in cluster g1 and 99.8% in cluster g2 are of DRG origin (1.5% of cluster g1+2 are cochlear cells), indicating that these two clusters are the DRG-specific satellite glia.

Differential gene expression analysis showed that many genes are expressed at higher levels in cluster g2 compared to cluster g1 (Figure S8), but that no genes are uniquely expressed in one cluster. Thus, rather than corresponding to two types of satellite glia, cells in clusters g1 and g2 seem to represent satellite glia at different developmental stages, with cluster g1 as the immature satellite glia. Indeed, a significantly greater number of cells from the E14 time point are present in cluster g1 (# cells: 4004) than in cluster g2 (# cells: 2405), and significantly more cells from the P14 time point are present in cluster g2 (# cells: 1980) than in cluster g1 (#cells: 1779). Chi-squared test of independence showed that significantly different number of cells from each time point are present in cluster g1 and g2 (X^2^ (3, *N* = 10,168) = 221.666, *p* = < .00001). This progressive shift from cluster g1 to g2 over developmental time is consistent with the hypothesis that cluster g1 represents an immature state of satellite glia, while cluster g2 represents a more mature satellite glia state, which is marked by higher expression of genes characteristic of DRG satellite glia. Thus, we will refer to clusters g1 and g2 collectively as DRG satellite glia.

Comparative analysis of genes expressed in all three satellite glia clusters identified shared expression of multiple genes. This satellite glia signature gene set included known satellite glia markers such as *Kcnj10/Kir4*.*1* and *Glul*, as well as *Bcan, Cxcr4, Fgfr1, Nrarp, Adamts5, Hmgcs1, Cst3, Atp1b2* and *Glast1* (Figure 5B). Interestingly, these genes are also characteristically expressed in astrocytes (Anlauf and Derouiche, 2013; Bachoo et al., 2004; Batiuk et al., 2017; Cali and Bezzi, 2010; Clarke et al., 2001; Demircan et al., 2013; Hu et al., 2019; Itoh et al., 2018; Shibata et al., 1997; Tong et al., 2014; Yamada et al., 1997), suggesting functional similarities between satellite glia and astrocytes, as highlighted recently (Avraham et al., 2020).

In stark contrast to the uniformity between Schwann cells from the two ganglia, SG and DRG satellite glia diverge significantly beyond this core set of signature genes. Although SG satellite glia can be recognized by the presence of known markers, *Glul/GS* and *Kcnj10/Kir4*.*1* (Figures 5A, 5B), these genes are not restricted to satellite glia in the literature or in our dataset (Chen and Zhao, 2014; Hanani, 2005; Liu et al., 2014). To identify a more selective marker for SG satellite glia, we performed differential gene expression analysis and identified *Igfbp6* (*Insulin Like Growth Factor Binding Protein 6*) (Figure 5A). As predicted from the scRNA-seq data, FISH by Hybridization chain reaction (HCR) method at P14 confirmed expression of *Igfbp6* only in SG satellite glia and not in DRG, SN or RB (n=10 fields each for SG & RB from 5 ears, 5 mice; n=16 DRG and 15 SN fields from 6 DRGs and 2 SN per animal, 3 mice) (Figure 5C). These data validate cluster g11 as SG satellite glia and also provides a unique marker for this glial subtype. Igfbp6 is a secreted O-linked glycoprotein that binds IGF-II (Bach, 2015) as well as the single-span membrane protein Prohibitin-2 (Phb2) (Fu et al., 2013). Although IGF-II is not enriched in SG satellite glia nor in SG neurons in our dataset and others (Petitpre et al., 2018) (Figure S9A), Phb2 is highly expressed in SG neurons (Shrestha et al., 2018) (Figure S9B). Thus, this glial-neuronal gene pair (Igfbp6-Phb2) may reflect a distinctive glial-neuronal communication between SG satellite glia and neurons.

DRG satellite glia, on the other hand, are distinguished by high levels of *Epas1* (*Endothelial PAS Domain Protein 1*), which is not expressed by SG satellite glia or by diverse Schwann cell types (Figure 5A). Validation by RNAscope confirmed unique expression of *Epas1* in DRG satellite glia, but not in the SN, cochlear SG or RB at P14 (n=8 fields each for SG & RB from 5 ears, 5 mice; n=13 DRG and 10 SN fields from 6 DRGs and 2 SN per animal, 6 mice) (Figure 5D). *Epas1* is expressed in all the stages that we profiled, indicating that its expression starts early and continues into postnatal life. Furthermore, *Epas1* was also defined as a top marker for the DRG GP cluster g9 and the DRG GPs found in the mixed cluster g4 (Figure 5A, heatmap in Figure 2E). Indeed, many of the top genes highly expressed in DRG satellite glia clusters are also highly expressed by cluster g9 DRG GPs (Figure S10A). Analogously, genes enriched in cells of cluster g9 compared to all other glia include genes expressed by both clusters of DRG satellite glia in addition to the proliferative gene set shared with other GP clusters (Figure S10B).

Together, these findings suggest that the DRG satellite glia retain a gene signature found in the DRG-specific GPs from which they derive, whereas this gene signature is lost in Schwann cells that also derive from these GPs.

### Myelination-related genes show differential expression in SG vs DRG satellite glia

Consistent with their known functional differences, comparison of SG and DRG satellite glia revealed that many genes distinctive for SG satellite glia are associated with myelination and myelin, including *Mbp, Mpz, Pmp22, Prx, Egr2, Gldn, Fa2h*, and *Gjb1/Cx32*. All of these genes are also highly expressed in the MSC cluster but are not in DRG satellite glia (Figure 4C, Figure S11A). Consistent with expression of this myelination program, most SG neuron cell bodies are myelinated in mice and other species (Hibino et al., 1999; Spoendlin, 1967; Toesca, 1996). This is in stark contrast with DRG neuronal cell bodies, which are not myelinated (Nascimento et al., 2018). To characterize differences in myelination further, we examined expression of Mbp protein and found clear expression in satellite glia within cochlea but not within DRG from P14 mice (n=2 fields each for SG & RB from 2 ears, 2 mice; n=6 DRG fields from 6 DRGs per animal, 3 mice) (Figure 6A). SG satellite glia also resemble MSCs in that they express genes such as *Arhgap19, Srgn, Fxyd6* and *Sema5a*, while DRG satellite glia do not (Figure S11A). Arhgap19 controls cytokinesis (David et al., 2014), Srgn is a proteoglycan, Fxyd6 is an ion transport regulator, and Sema5a is a member of the semaphorin family implicated in axon guidance (Kantor et al., 2004). We suspect these genes may function in the morphologic changes required during myelination in SG satellite glia as well as in MSCs. However, some genes normally associated with axon myelination, such as *Pllp, Ncmap, Pou3f1* and *Mag*, are not expressed in SG satellite glia, which instead myelinate the cell body (Figure 6B). Thus, the myelination process and/or the myelin structure may be different in SG satellite glia than in MSCs.

**Figure 6:**
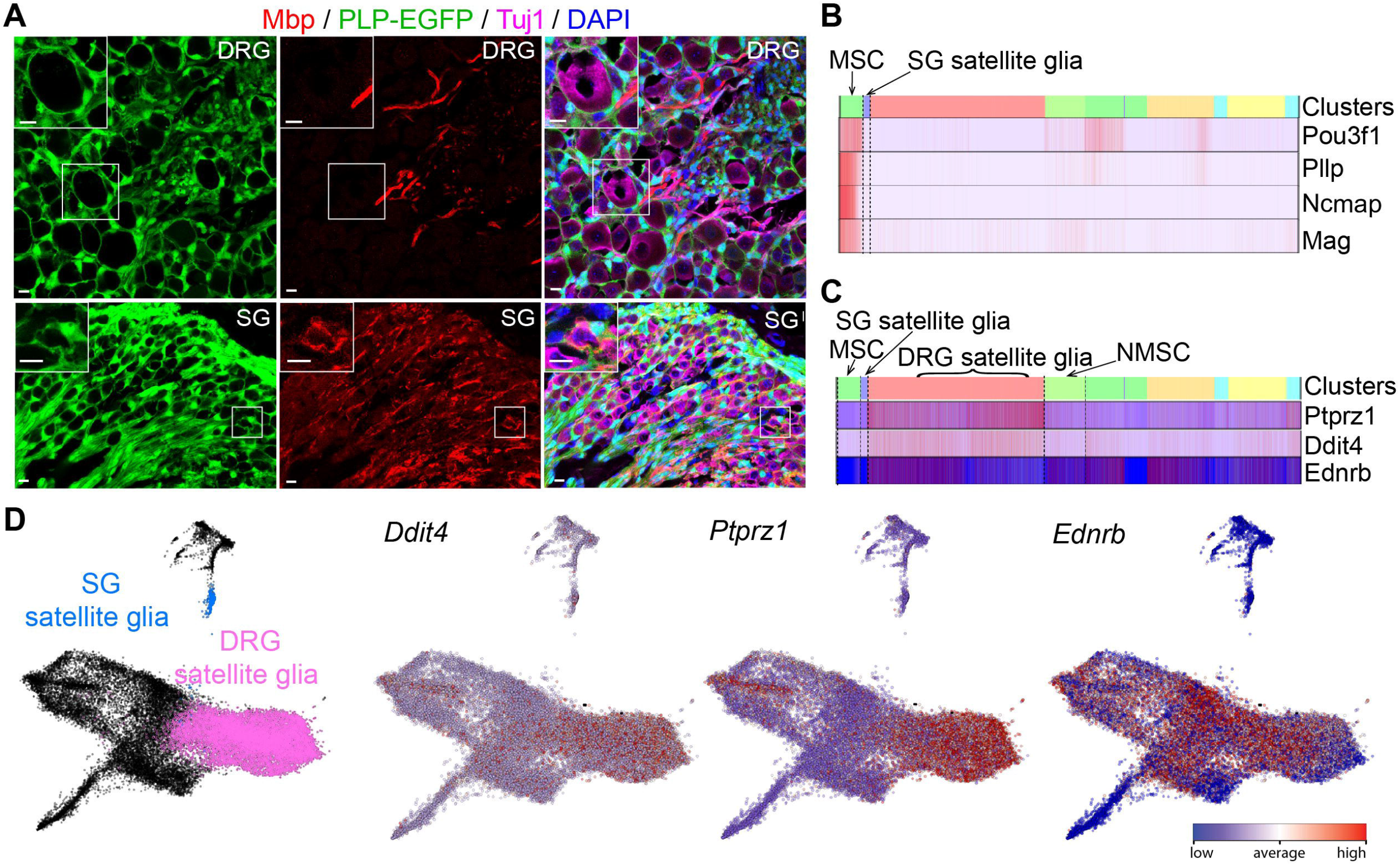
Myelination-related genes show differential expression in DRG vs SG satellite glia (A) P14 DRG and SG tissue sections label glia with PLP-EGFP in green, myelin marker Mbp in red and neurons with Tuj1 in magenta. Mbp is present in many satellite glia from SG, but not from DRG. Scale bars: 10um (B) Heatmaps of myelin related genes expressed in MSCs but not in SG satellite glia. (C, D) Myelination inhibitors Ptprz1, Ddit4 and Ednrb are enriched in DRG satellite glia compared to SG satellite glia (heatmaps (C) and Joint embeddings (largeVis) (D)) and enriched in NMSCs compared to MSCs (heatmaps in C).

Concordant with the lack of myelin-related factors in the DRG satellite glia, our data suggest that myelination is actively inhibited in DRG satellite glia. DRG satellite glia, unlike SG satellite glia, express high levels of several genes reported to inhibit myelination including *Ddit4, Ptprz1* and *Ednrb* (Figure 6C, D). Moreover, these same genes are expressed by more cells in the NMSC cluster than in the MSC cluster, suggesting that myelination is also inhibited in NMSCs (Figure 6C). Ddit4 is a negative modulator of myelination that activates the tuberous sclerosis complex TSC1/2 and inhibits the actions of the mTOR pathway in promoting myelination (Noseda et al., 2013). Ptprz1 acts as a negative regulator of oligodendrocyte differentiation and myelination (Kuboyama et al., 2012). Endothelins and the receptor Ednrb inhibit Schwann cell maturation and support Schwann cell precursor survival (Brennan et al., 2000; Quintes et al., 2016). Together, these data suggest that myelination is continuously and selectively inhibited in DRG satellite glia, preventing the myelination process that occurs in SG satellite glia and in MSCs.

SG and DRG satellite glia differ in other respects that may relate to the requirements of the associated sensory neurons. Several genes enriched in SG satellite glia compared to DRG satellite glia may have unique developmental or functional roles in the cochlea (Figure S11A). For example, Slc6a15, which functions in reuptake of neurotransmitters (Drgonova et al., 2007), and Syt4, which is essential for glutamate release from astrocytes (Zhang et al., 2004), could allow SG satellite glia to influence neurotransmission by the SG neurons. Genes that are instead enriched in DRG satellite glia include several implicated in synapse formation and function (*Sparc, Sparcl1, Tspan7*), suggesting distinct mechanisms for modulation of DRG neuron firing (Figure S11B). DRG satellite glia also show enriched expression of a set of genes related to lipid and energy metabolism (Figure S11B), consistent with the recent demonstration that a *de novo* fatty acid synthesis protein, Fasn (Fatty Acid Synthase), acts in DRG satellite glia to promote axon regeneration (Avraham et al., 2020). Consistent with that finding, our dataset showed strong expression of *Fasn* in DRG satellite glia and in other peripheral glial types (data not shown). DRG satellite glia also express high levels of *Apoe*, which encodes a plasma protein that is an important determinant of lipid transport and metabolism (Figure S11B) and is expressed both in CNS astrocytes and in nonmyelinating peripheral glia (Boyles et al., 1985). ApoE levels are reported to increase after injury (Boyles et al., 1990) and ApoE itself, like Fasn, may promote axon regeneration (Li et al., 2010). Taken together, these data show that satellite glia in the two ganglia exhibit clear differences in gene expression and our dataset indicates new avenues to investigate unique functions of these cells.

### Placode-derived supporting cells and NC-derived glia share a core glial signature gene set

Although satellite glia and Schwann cells are the most prevalent types of glia in the PNS, the cochlea also houses specialized cells that express classic glial markers. These cochlear supporting cells surround and support the hair cells, the sensory receptor cells that initiate and relay sound information to SG neurons (Figure 7A). Supporting cells maintain ion homeostasis, clear neurotransmitters, promote synapse formation, and phagocytose debris, functions that are associated with astrocytes and that are also observed in invertebrate glia (Freeman and Rowitch, 2013; Wan et al., 2013). In contrast to NC-derived peripheral glia, supporting cells originate in the otic placode, an epidermal thickening that invaginates to form the entire inner ear, including its sensory organs and neurons. Despite this distinct embryonic origin, supporting cells also express PLP-EGFP (Figure S1B) and were thus present in cochlear samples at all three ages. To learn more about shared and unique features of peripheral glia, we compared these unusual glia-like cells to NC-derived peripheral glia.

**Figure 7:**
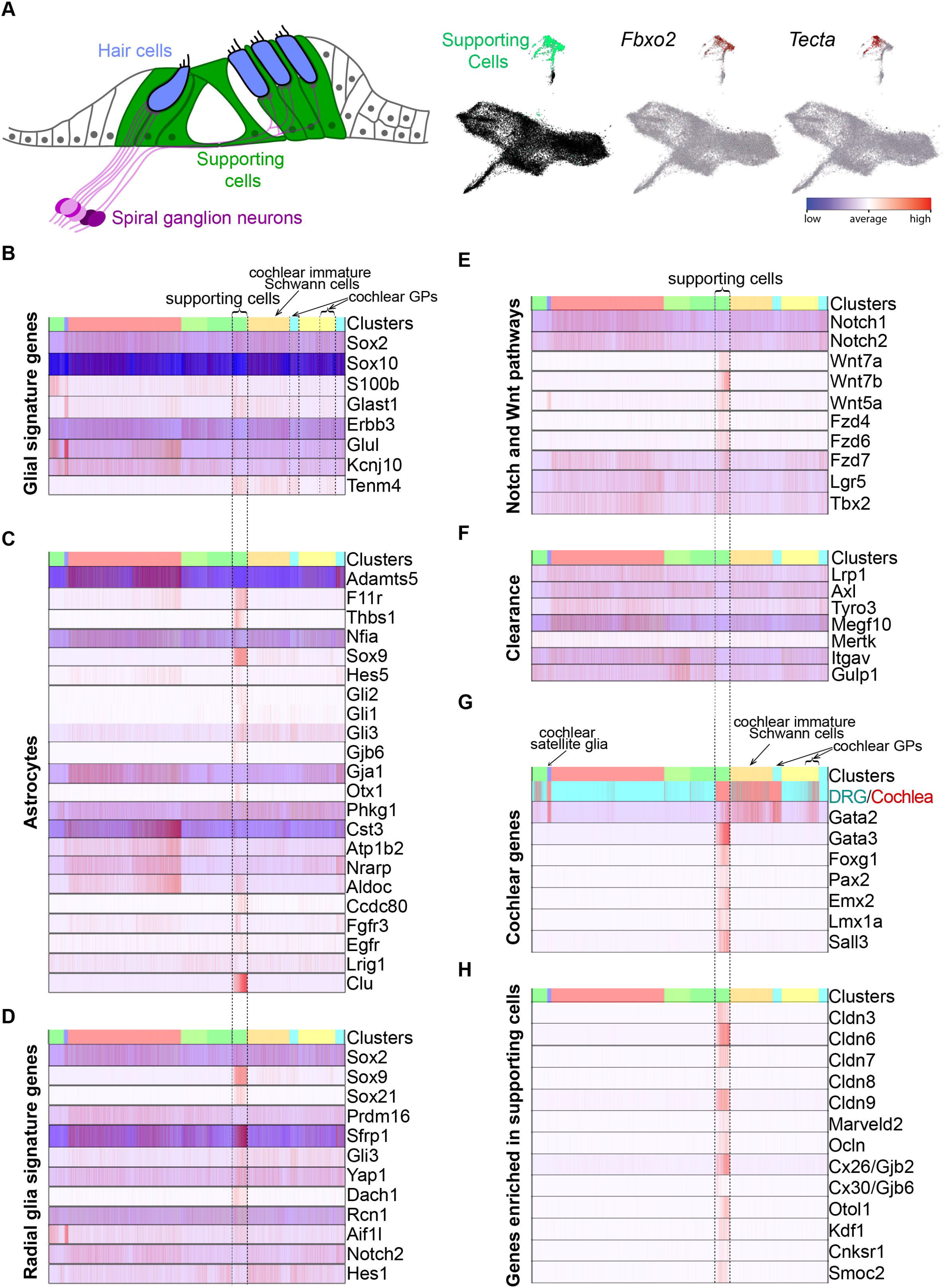
Placode-derived supporting cells and NC-derived glia share a core glial signature gene set. (A) Schematic describes supporting cells. Joint embeddings (largeVis) show supporting cell cluster in green and expression of Fbxo2 and Tecta. (B-H) Heatmap of genes that are (B) signature of glia (C) astrocytic (D) signature of radial glia (E) associated with Notch and Wnt pathways (F) cochlear (G) regulating clearance (H) enriched in supporting cells.

Conos analysis revealed that supporting cells are transcriptionally distinct from the other peripheral glia, yet still express many genes associated with the glial identity. A single cluster of supporting cells (cluster g8) was identified based on the expression of the known markers *Fbxo2* and *Tecta* (Hartman et al., 2015; Kim et al., 2019) (Figure 7A). Although there are several different types of adult supporting cells (Wan et al., 2013), our analysis revealed a single cluster, possibly due to the underrepresentation of P14 cells relative to those from embryonic stages. Consistent with their distinct morphologies, functions, and embryonic origins, this supporting cell cluster is positioned separately from most other peripheral glia clusters. Nonetheless, supporting cells and peripheral glia express many shared genes, including well-established glial genes such as *Sox2, Sox10, S100b, Glast1, Erbb3*, and *Kcnj10/Kir4*.*1*, consistent with earlier studies (Figure 7B) (Boettger et al., 2002; Boettger et al., 2003; Furness and Lehre, 1997; Hume et al., 2007; Pak and Slepecky, 1995; Wan and Corfas, 2017; Watanabe et al., 2000; Zhang et al., 2014). In addition to these familiar genes known to be important for glial development and function, we found Teneurin-4 (Tenm4), a regulator of oligodendrocyte differentiation and myelination (Suzuki et al., 2012), to be expressed in supporting cells as well as in cochlear GPs and immature Schwann cells (Figure 7B). Together these findings confirm the glial nature of the supporting cells and help define a core set of glial signature genes.

Many of the characteristic glial genes expressed by supporting cells and by diverse satellite glia are also typically expressed by astrocytes, a major CNS glial cell type. This gene cohort includes *Glast1, Kcnj10/Kir4*.*1, Glul, Erbb3*, and *S100b* (Figure 7B), as well as several genes encoding proteins related to modulation of the ECM, such as the ECM degrading enzyme Adamts5 and the cell adhesion molecule F11r/JAM-A (Figure 7C) (Demircan et al., 2013; Horng et al., 2017). In addition, the gene encoding the matricellular protein Thbs1, which is secreted from astrocytes and modulate synapses, is enriched in supporting cells compared to other glia, suggesting that supporting cells may have similar effects on synapse formation or function (Figure 7C) (Christopherson et al., 2005). Supporting cells and satellite glia also exhibit shared expression of genes encoding astrocytic transcription factors Nfia, Sox9, and Hes5 (Figure 7C) (Deneen et al., 2006; Hasel et al., 2017; Kang et al., 2012). In the CNS, Sox9 induces expression of NFIA, which influences the onset of gliogenesis and astrocyte differentiation.

Subsequently, Sox9 and NFIA can interact to coregulate a defined geneset during gliogenesis. Although *Nfia* is expressed in all peripheral glia, Sox9 is only expressed in cochlear GPs and supporting cells, suggesting that NFIA acts with a coregulator other than Sox9 in the somatosensory system. Lastly, supporting cells express some of the same connexins as astrocytes: *Cx30/Gjb6* is enriched in supporting cells and *Cx43/Gja1* is shared with both supporting cells and satellite glia (Figure 7C) (Rouach et al., 2008; Stogsdill and Eroglu, 2017). Thus, across the central and peripheral nervous systems, this shared geneset may enable diverse glial cell types to shape the neuronal milieu and facilitate intercellular communication.

In addition to the similarities shared by supporting cells and astrocytes, supporting cells also share features with radial glia, the multi-potent CNS stem cells that produce astrocytes as well as neurons, oligodendrocytes, and ependymal cells. Like radial glia, supporting cells are radial and span the depth of the nascent sensory epithelium (Figure 7A). Additionally, supporting cells show stem-cell like properties and can initially serve as progenitors for hair cells, a property that is gradually lost in older mammals but retained in adult amphibians and birds (Stone and Cotanche, 2007). These gross similarities are reflected in the transcriptome: supporting cells express “radial glia signature genes” associated with stem cell properties, such as *Sox2, Sox9*, and *Sox21* (Figure 7D) (Lui et al., 2014). Multiple components of the Notch and Wnt signaling pathways, which control stem cell behavior throughout the body are expressed in supporting cells, and in other peripheral glia (Figure 7E). The Wnt pathway-associated receptor Lgr5 is a well-established stem cell marker that is expressed by supporting cells (Chai et al., 2012; Shi et al., 2012) and, as shown here, by GPs, NMSCs, and DRG satellite glia (Figure 7E). Furthermore, all of these peripheral glia express the gene for the transcription factor Tbx2 (Figure 7E), which functions downstream of canonical Wnt signaling (Aydogdu et al., 2018). These data raise the possibility that Tbx2 and Lgr5 cooperate to maintain more plastic states in multiple types of peripheral glia.

Supporting cells and peripheral glia also share with astrocytes another crucial role in maintenance of the nervous system: along with macrophages they participate in clearance of apoptotic cells and cellular debris. Schwann cells clear axon and myelin debris after injury (Brosius Lutz et al., 2017; Gomez-Sanchez et al., 2015) and satellite glia clear apoptotic cells (Wu et al., 2009). In response to several insults, hair cells undergo cell death and are removed by supporting cells (Abrashkin et al., 2006; Anttonen et al., 2014), in addition to macrophages. Such removal is crucial for prompt sealing of the epithelium by supporting cells so that the barrier between high-potassium endolymph and low-potassium interstitial perilymph is maintained (Oesterle, 2013). We found that supporting cells and/or peripheral glia express several genes that encode proteins implicated in phagocytosis and clearance, including Clu (Cunin et al., 2016) (Figure 7C), Lrp1 (Liu et al., 2017), Axl and Tyro3 (Lemke and Burstyn-Cohen, 2010) (Figure 7F). *Clu* is strongly enriched in supporting cells, indicating an active role in clearance. Future studies will uncover when these genes are utilized by glial cells to eliminate cellular debris.

Among the peripheral glia, SG satellite glia are most similar to supporting cells at the transcriptional level. The proximity of the supporting cell and SG satellite glia clusters suggests that NC-derived and otic placode-derived progenitors are similarly shaped by signals in the environment that impart a cochlear identity. In support of this idea, supporting cells, SG satellite glia, and cochlear GPs all express the gene encoding the transcription factor Gata2 (Figure 7G), which is also expressed by efferent auditory neurons (Lillevali et al., 2004; Nardelli et al., 1999; Pata et al., 1999). In contrast, cochlear Schwann cells, which are transcriptionally indistinguishable from DRG Schwann cells, do not show high levels of *Gata2* expression. This raises the possibility that convergent development of Schwann cells involves downregulation of Gata2 and hence of the auditory identity.

Several other transcription factors enriched in supporting cells compared to other peripheral glia also play prominent roles in inner ear development. For instance, supporting cells express *Pax2, Gata3*, and *Foxg1*, all of which are expressed in the early otic placode and required for multiple features of inner ear development (Figure 7G) (Appler et al., 2013; Burton et al., 2004; Pauley et al., 2006). These genes are expressed by several other cochlear cell types, such as hair cells and neurons, presumably reflecting a common placodal origin. Supporting cells are also distinguished from other peripheral glia by the presence of *Emx2* and *Lmx1a*, both of which are also required for inner ear patterning and morphogenesis (Holley et al., 2010; Nichols et al., 2008), and by *Sall3*, whose role in the cochlea has not been defined (Figure 7G) (Parrish et al., 2004; Yang et al., 2020).

Despite their gross similarity to other glia, supporting cells ultimately take on distinct functional properties that are reflected by their gene expression profiles. For instance, whereas Schwann cells express genes needed for myelination, supporting cells play crucial roles in providing structure to the organ of Corti and insulating hair cells. Accordingly, supporting cells uniquely express several genes that encode tight junction proteins, including multiple claudins (3, 6-9), MARVELD2, and Ocln (Figure 7H). Given this tight epithelial structure, gap junctions may offer a critical route of intercellular communication, reflected in enriched expression of the gap junction proteins Cx30/Gjb6 and Cx26/Gjb2 (Figure 7H), (Lautermann et al., 1998). Mutations in *Gjb2* and *6* in *DFNB1* account for an estimated 50% of inherited non-syndromic deafness in a variety of populations (Alford et al., 2014), whereas *MARVELD2* is mutated in *DFNB49* (Najmabadi and Kahrizi, 2014), further emphasizing the crucial contributions of supporting cells to cochlear function.

## Discussion

The remarkable variety of neurons responsible for different sensory modalities demands additional diversity among the peripheral glia associated with them. At present, little is known about the molecular basis for these differences or how they arise during development. Here, we present a comprehensive analysis of transcriptomes for the peripheral glial subtypes associated with two different sensory ganglia, DRG and SG. Using single cell RNA-seq technology and the Conos algorithm, we analyzed gene expression of these diverse glial cells across multiple developmental stages and identified a core set of genes that are common to glia throughout the peripheral and central nervous systems. While multiple types of glial cells express this core glial gene program, our data suggest that glial cells, from early GPs to mature glial cells, express distinct sets of genes and follow unique developmental trajectories depending on their rostro-caudal position and on the nature of the sensory system within which they function.

Despite these developmental differences, neither MSCs nor NMSCs exhibit gene expression profiles that are specialized for location or sensory modality, offering a new example of convergent differentiation. Our data also revealed better markers to label and manipulate developing and mature glial subtypes and identified gene expression features suggestive of a novel type of interaction between satellite glia and neurons in the cochlea. Together, our results provide a comprehensive view of the development and function of diverse peripheral glia with both shared and unique properties, introduce improved markers for glial subtypes, and suggest new hypotheses for how specializations among peripheral glia influence sensory function. These data will serve as a beneficial resource for other scientists seeking to investigate the origin and impact of glial diversity for PNS function and dysfunction.

Our analyses revealed that somatosensory and cochlear GPs exhibit significant differences in their molecular nature and developmental trajectories and these differences reflect their origins and their locations. Cochlear GPs develop from cranial NCCs, whereas lumbar DRGs arise from trunk NCCs. Although cranial and trunk NCCs are largely similar, they also display axial-specific differences in gene expression (Soldatov et al., 2019). Some of these axial-specific genes are also expressed in the cochlear and DRG GPs, suggesting that early differences in the NCC are maintained in the GPs. For instance, we observed continued expression of the cranial NCC genes *Dlx2* and *Sox9* in cochlear GPs. Beyond these origin-related differences, the GPs migrate along different routes and mature within different tissues, so local interactions in these distinct environments may contribute to some of the region-specific programs that we observe, such as enriched expression of Epas1 in lumbar DRG GPs compared to cochlear GPs. Epas1 is a transcription factor that promotes stem cell renewal in other systems (Rankin et al., 2008) and could enable maintenance of this precursor population in adult DRGs. In support of this possibility, we find that GPs are present at early stages in both ganglia but are almost absent in the adult cochlea, whereas much higher numbers of GPs persist in adult lumbar DRG+SN. Cochlear GPs, on the other hand, are poised to directly affect nearby SG neurons through production of the neuropeptide Npy. The availability of distinct gene expression profiles for DRG and cochlear GPs provides tools for developing new models for peripheral glial tumors such as cochleovestibular schwannomas and peripheral nerve sheath tumors.

Although GPs from the two ganglia have distinct properties, we found evidence for convergent differentiation during the generation of Schwann cell subtypes (NMSCs and MSCs). In contrast to the GPs, NMSCs and MSCs from these ganglia are clustered together, reflecting similar gene expression profiles. Thus, specialized GPs produce mature Schwann cells without obvious tissue-specific differences. In other systems, distinct progenitors can also give rise to molecularly similar cell types (Chan et al., 2019; Liu et al., 2019b; Wagner et al., 2018). Convergent differentiation may be a common feature during the development of myelinating cells, as oligodendrocyte precursor cells (OPCs) from different origins also converge into a transcriptionally uniform pool of OPCs at postnatal stages (Marques et al., 2016).

Convergent differentiation for both Schwann cells and oligodendrocytes suggests there may be uniformity in complex gene programs that enable axon wrapping or myelination. Mechanistically, convergence may be driven by transcription factors such as Pou3f1, Sox10 and Brn2, which coregulate Egr2 and drive the transition from promyelinating to myelinating Schwann cell, including activation of genes encoding myelin components such as Mbp, Mpz, Cx32 (Svaren and Meijer, 2008). In our dataset, *Pou3f1* and *Egr2* are strongly expressed in MSCs compared to other mature glia, as expected, and presumably this drives all MSCs to express this myelination program, despite their distinctive developmental origins. Despite the molecular similarity between Schwann cells from the auditory and somatosensory systems, we found that there is a larger proportion of MSCs in the cochlea compared to SN, reflecting the need for high velocity transmission in the auditory system. This difference in NMSC:MSC proportion is likely due to differences in the composition of neurons in each system, given the known effect of axon caliber on the NMSC/MSC fate choice (Nave and Werner, 2014).

Unlike Schwann cells, satellite glia from DRG and cochlear tissues retain tissue-specific differences. Even though these satellite glia share expression of many core genes, they clustered separately and showed major differences in transcriptional profiles. Cochlear signature genes such as *Gata2*, a transcription factor, are expressed in auditory GPs and satellite glia but not in somatosensory glia. Additional differences among the satellite glia may be driven by the sensory systems in which they function, as the associated neurons convey distinct sensory information (sound versus pain, temperature, and touch) and are derived from distinct precursors (placode versus NC). Consistent with the fact that satellite glia are specialized according to the needs of the neurons, we find that SG satellite glia express many myelination-promoting genes (e.g. *Mpz*), whereas DRG satellite glia express genes that inhibit myelination (e.g. *Ptprz1*). The need for myelination of SG neuronal cell bodies may be due to their morphology. SG neurons are bipolar, and so the action potential must traverse the large neuronal soma, which has a high capacitance. As there is a need for fast transmission in the auditory system, myelination of the entire soma of SG neurons, in addition to myelination of the axons, may prevent delay or drop off of action potential. In contrast, DRG neurons are pseudounipolar, a T-shaped morphology that enables the action potential to bypass the stem axon and soma, reducing the risk of propagation failure due to capacitative loading (Luscher et al., 1994). As DRG satellite glia express multiple genes that inhibit myelination, we postulate that there is an active process to maintain DRG satellite glia in an unmyelinating state. In the CNS, somatodendritic expression of cell surface adhesion proteins in neurons can inhibit myelination by oligodendrocytes (Redmond et al., 2016); similarly, DRG neurons may signal DRG satellite glia to inhibit myelination.

In addition to these major differences in myelin-related genes, SG and DRG satellite glia each express unique sets of genes that provide insights into how they diverge developmentally and indicate their distinct functions in the mature PNS. For example, we found that *Epas1*, which encodes a hypoxia-inducible transcription factor, is expressed in DRG GPs and then maintained in mature DRG satellite glia. Epas1 promotes stemness in other systems (Zhu et al., 2016), and could therefore contribute to the maintained proliferative potential of GPs in DRGs. Additionally, our data suggest that Epas1 may also indirectly prevent myelination via regulation of the myelination-inhibitor Ddit4. *Ddit4* is also expressed in DRG satellite glia, is hypoxia-inducible, and its promoter/enhancer has at least one binding site for Epas1 (Geiger et al., 2011). SG satellite glia, on the other hand, show enriched expression of *Igfbp6*, which encodes a secreted glycoprotein that can bind the membrane protein Phb2 (Fu et al., 2013). Phb2 is highly expressed in SG neurons in our dataset and others (Shrestha et al., 2018), suggesting that Igfbp6-Phb2 interactions may enable direct communication between SG satellite glia and neurons. Igfbp6 is also secreted from astrocytes and can inhibit neuronal differentiation *in vitro* (Barkho et al., 2006), but it is not known whether Igfbp6 produced by satellite glia exerts similar effects on SG neurons. Indeed, our data highlight the potential for many differences among satellite glia that remain to be explored, with differential expression of multiple genes related to lipid/energy metabolism, synaptic function, proliferation, ECM/adhesion, transporters/channels and various signaling pathway components. Thus, even though satellite glia in different sensory environments share a common ability to support neurons, the nature of this support is shaped to meet the local functional needs of each system. In the future, profiling satellite glia from other sensory ganglia may reveal distinctive cohorts of genes based on the needs of the sensory neurons with which they are associated.

As well as identifying the genetic potential for system-specific specializations, our data highlight broader commonalities of glia across the nervous system, regardless of embryonic origin. This is particularly apparent in the supporting cells, which express many genes typical of satellite glial cells, even though supporting cells develop from the otic placode, not NCCs. Many of these core glial signature genes are also shared with CNS radial glia and astrocytes. This shared astrocyte/satellite/supporting cell geneset includes components critical for synaptic function and homeostasis, as well as genes needed for clearance of apoptotic cells, a crucial role of glia in homeostatic maintenance. The common expression of these genesets in three morphologically distinct types of glia that arise from three distinct precursors suggests that a core glial function is the maintenance of electrically active cells, both by modulating synaptic transmission and by clearing debris from their environment. Notably, SG satellite glia and supporting cells are fairly similar to each other at the transcriptional level, even though they differ in their cell of origin, play distinct roles in the cochlea, and associate with vastly different kinds of sensory cells, with SG satellite glia enveloping SG neurons and supporting cells surrounding hair cells. The proximity of the supporting cell and SG satellite glia clusters, despite these differences, suggests that environmental signals dictate a cochlear identity to NC-derived and otic placode-derived progenitors. The transcription factor Gata2, which is strongly expressed in cochlear GPs, satellite glia and supporting cells, may regulate expression of multiple genes that confer common cochlear characteristics.

Taken together the data here provide new understanding of developing peripheral glial cells and supply a resource for future studies of the peripheral sensory system. In addition to their physiological roles in the development and function of a complex nervous system, peripheral glia have critical roles in pathological processes of neurodegeneration, inflammation, injury, tumor formation, and pain. This new understanding of peripheral glia will therefore facilitate future studies on the biology of sensory transduction and on human diseases including hearing loss, neuropathy, nerve injury, Schwannomas and chronic pain.

## ACKNOWLEDGEMENTS

We thank Drs. Lucas Cheadle and Mark Aurel Nagy for help with sequencing and library quantification; Kyomi Igarashi for help with the Indrop platform and library preparation; Dr. Qiufu Ma (DFCI) for sharing *Npy*^*Cre*^;*Ai14* mice; Drs. Lucas Cheadle and Yihang Li for helpful comments and editing. We thank the Dana-Farber Flow Cytometry Jimmy Fund Core for help with FACS analysis and sorting, and the Single Cell Core Facility of the ICCB-Longwood Screening Facility at Harvard Medical School for Indrop run. This work was supported by The Pussycat Foundation Helen Gurley Brown Presidential Initiative (to O.E.T.Y.), NIH/NINDS RO1 NS050674 (to R.A.S.), NIH T32AG000222 (to O.E.T.Y.), NIDCD RO1 DC009223 (to L.V.G.).

## Notes

### Competing Interest Statement

The authors have declared no competing interest.

