## Supplementary figures and images for "Diversity of developing peripheral glia revealed by single cell RNA sequencing"

### Figure S1.jpg

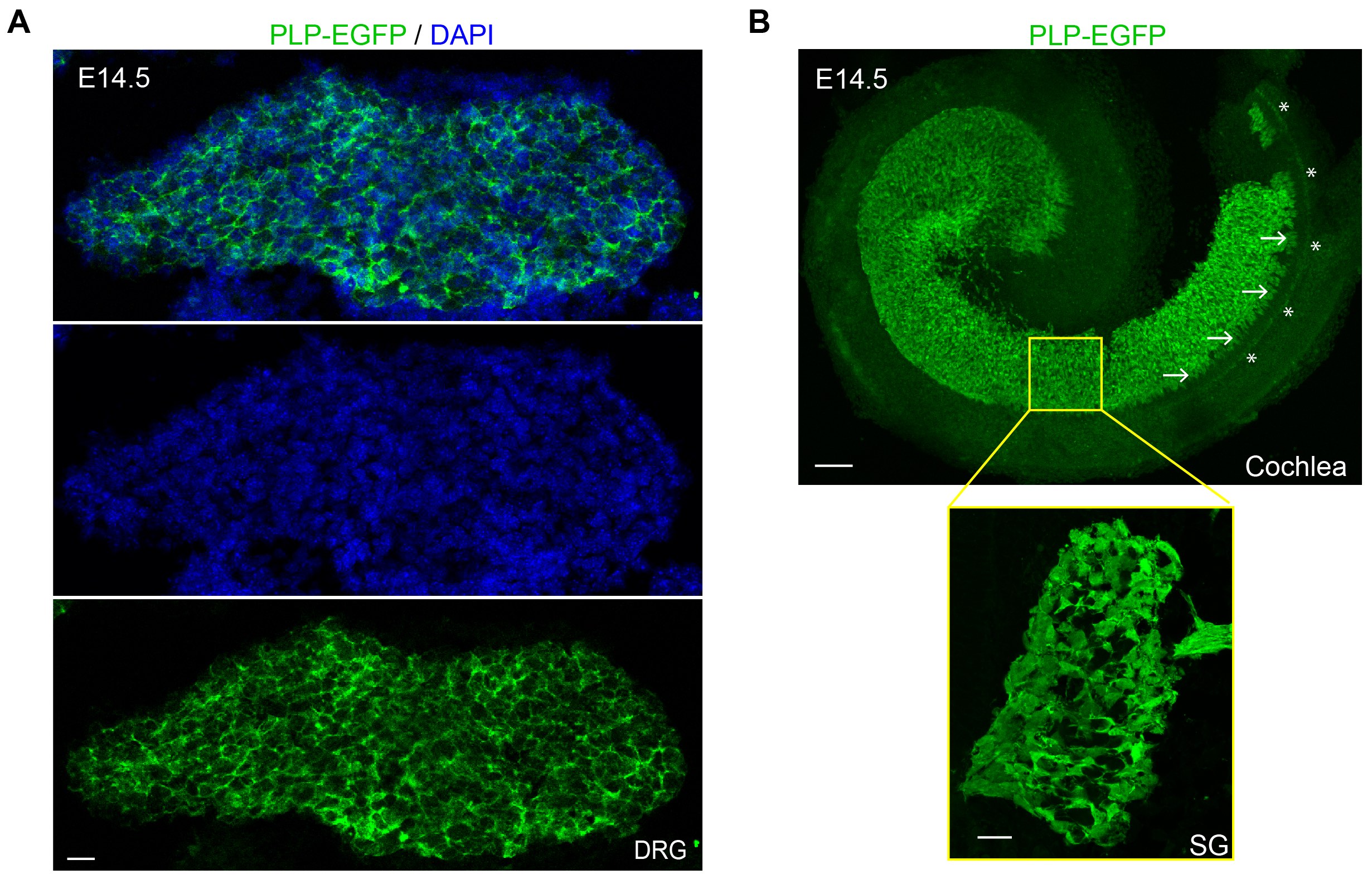

### Figure S2.tiff

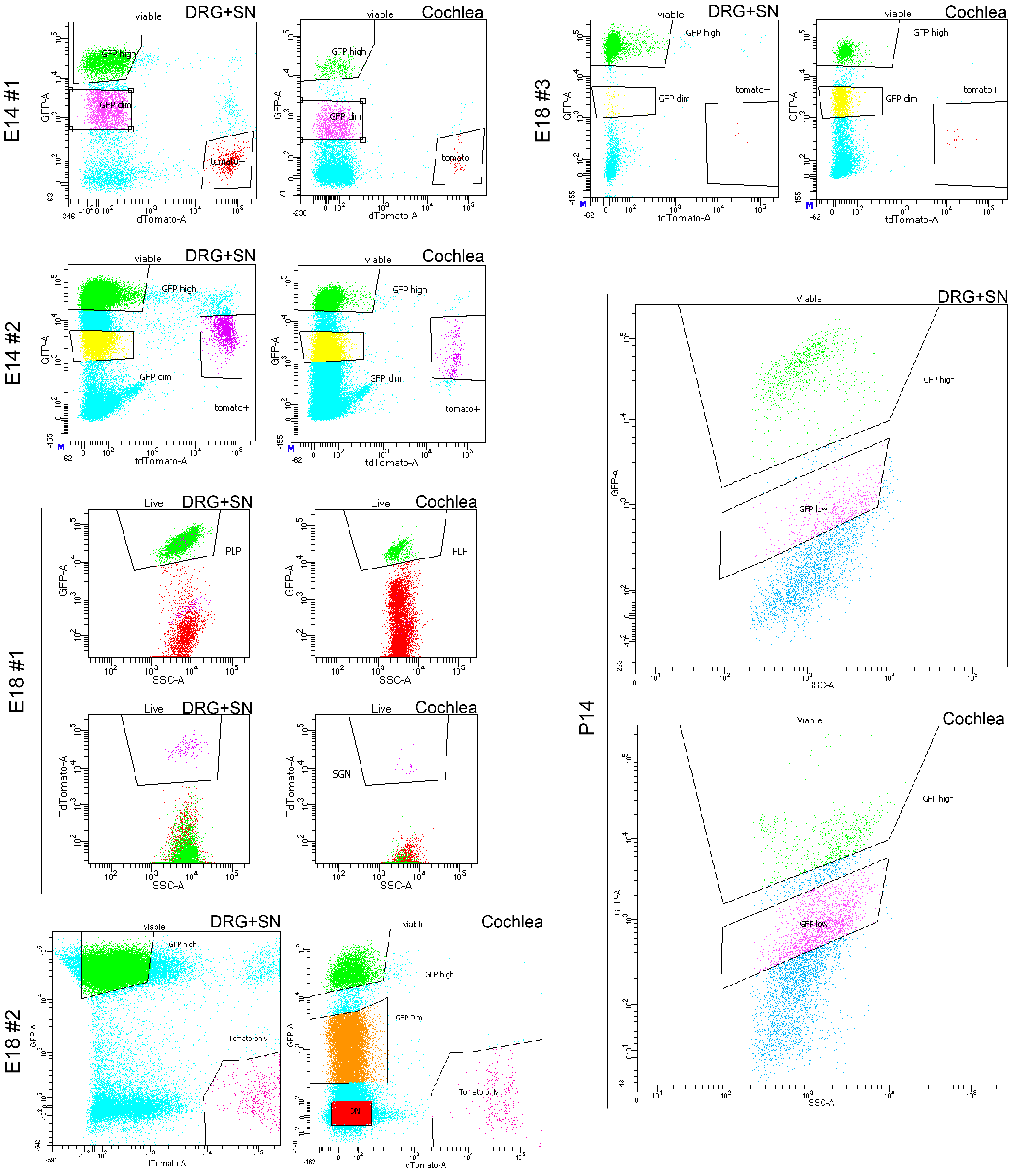

### Figure S3.jpg

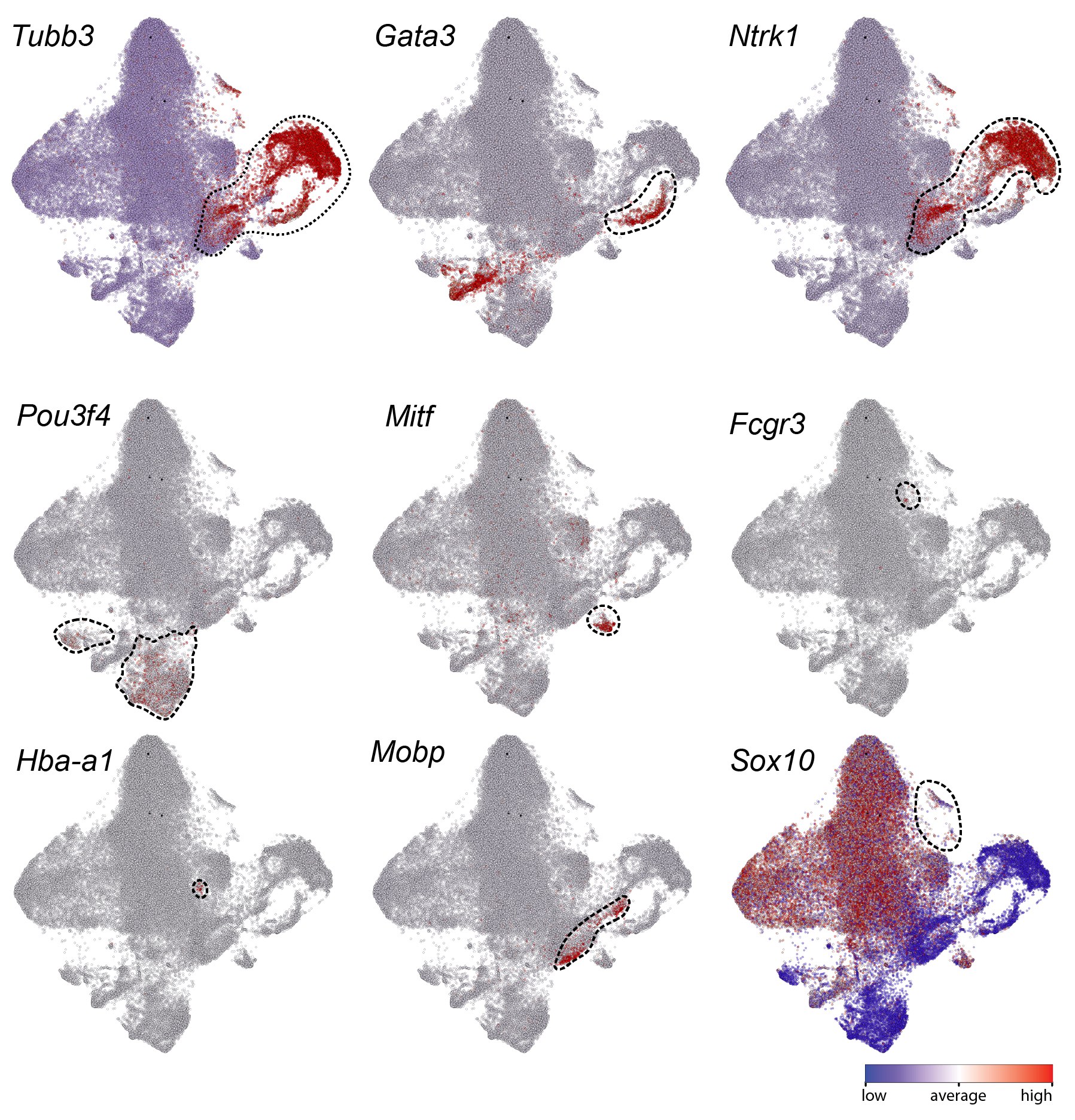

### Figure S4.jpg

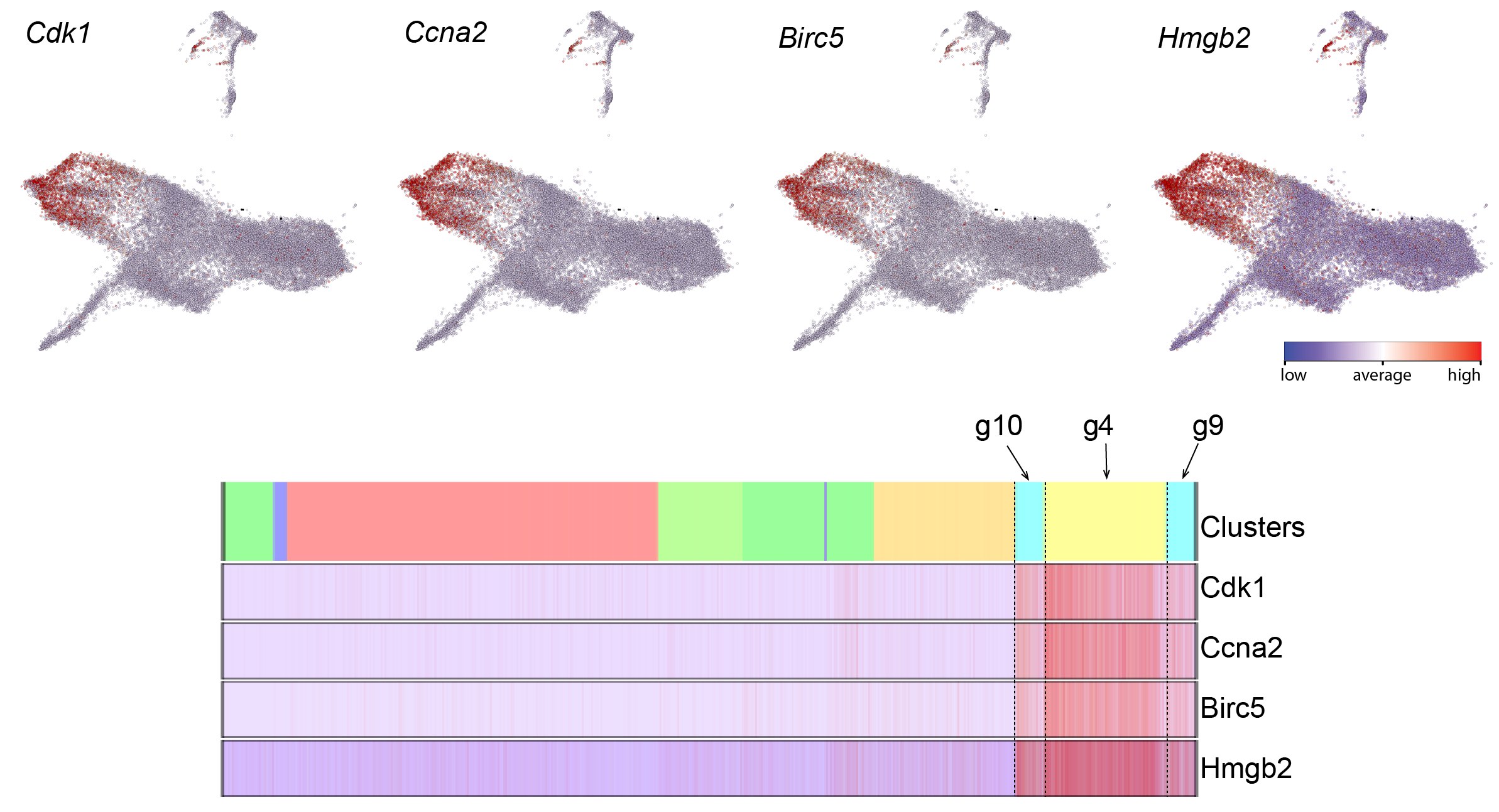

### Figure S5.jpg

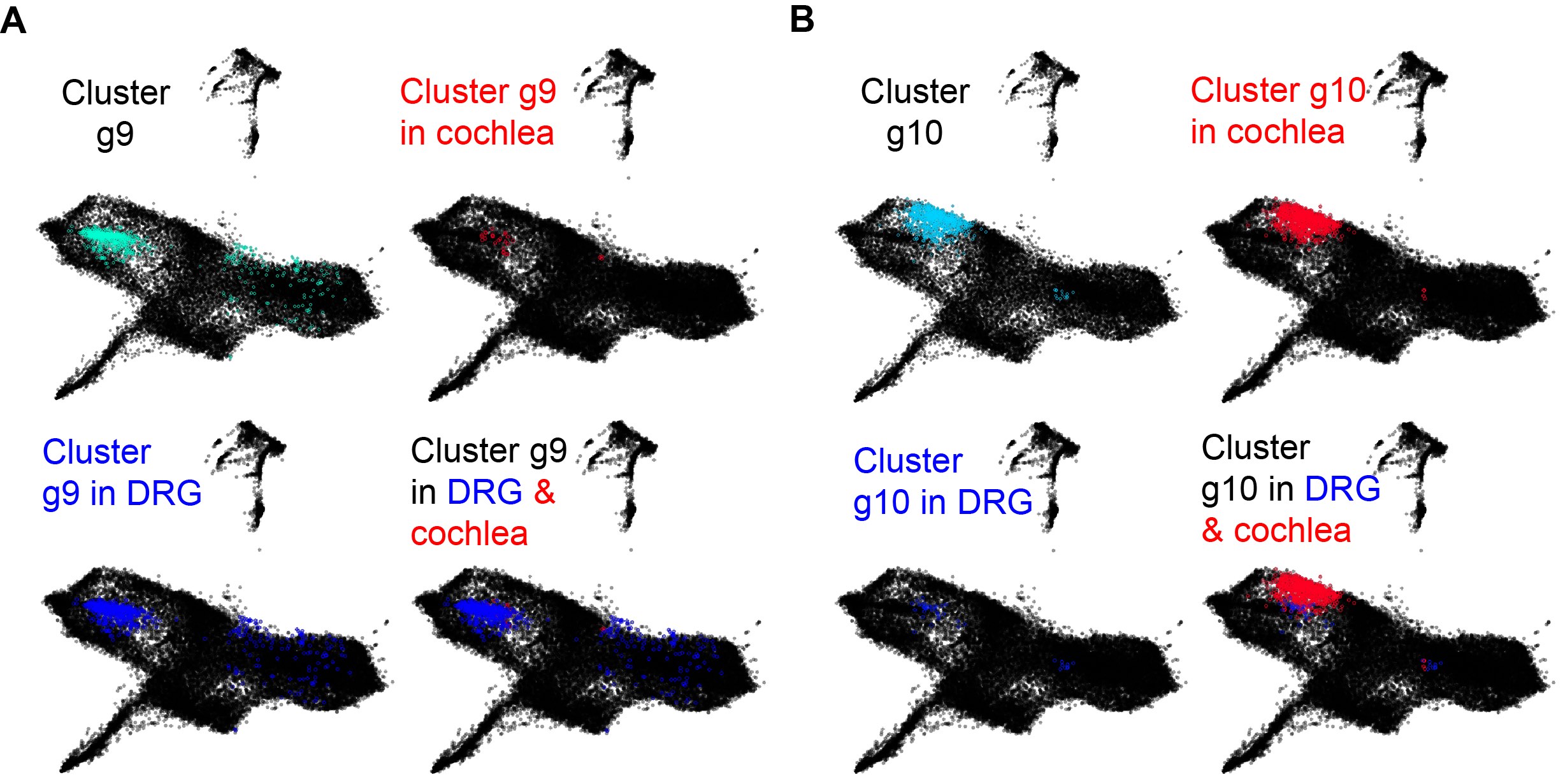

### Figure S6.jpg

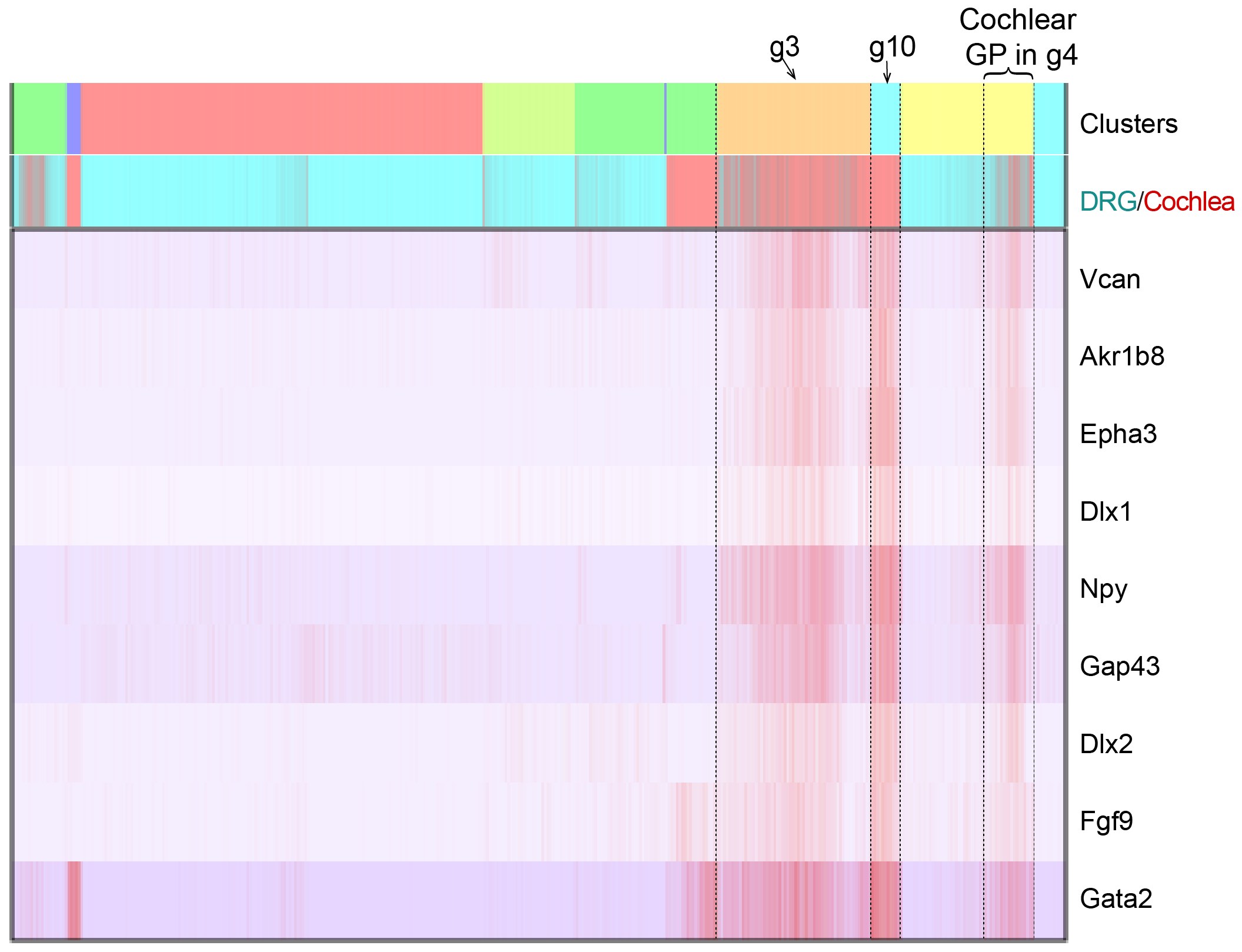

### Figure S7.jpg

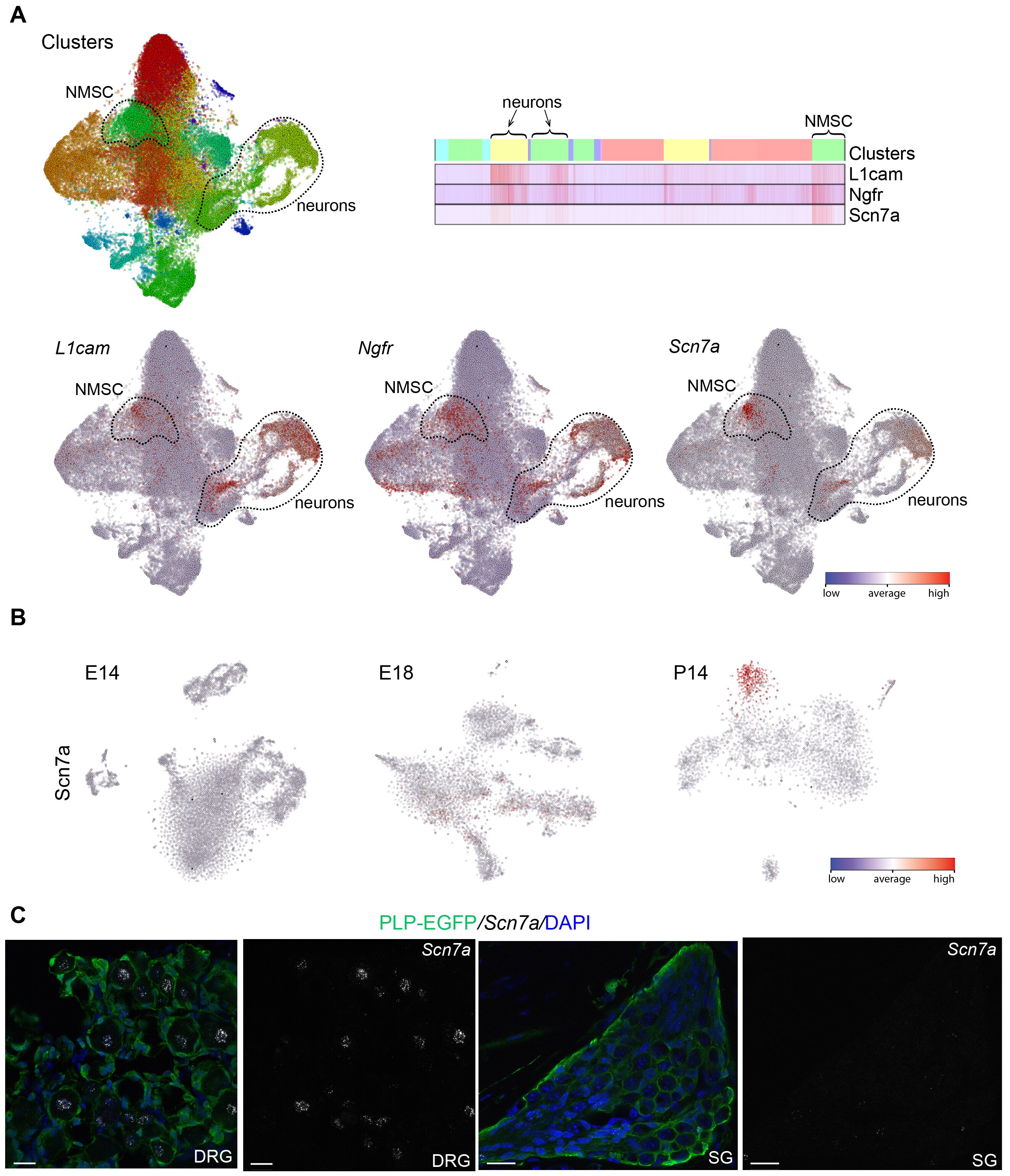

### Figure S8.jpg

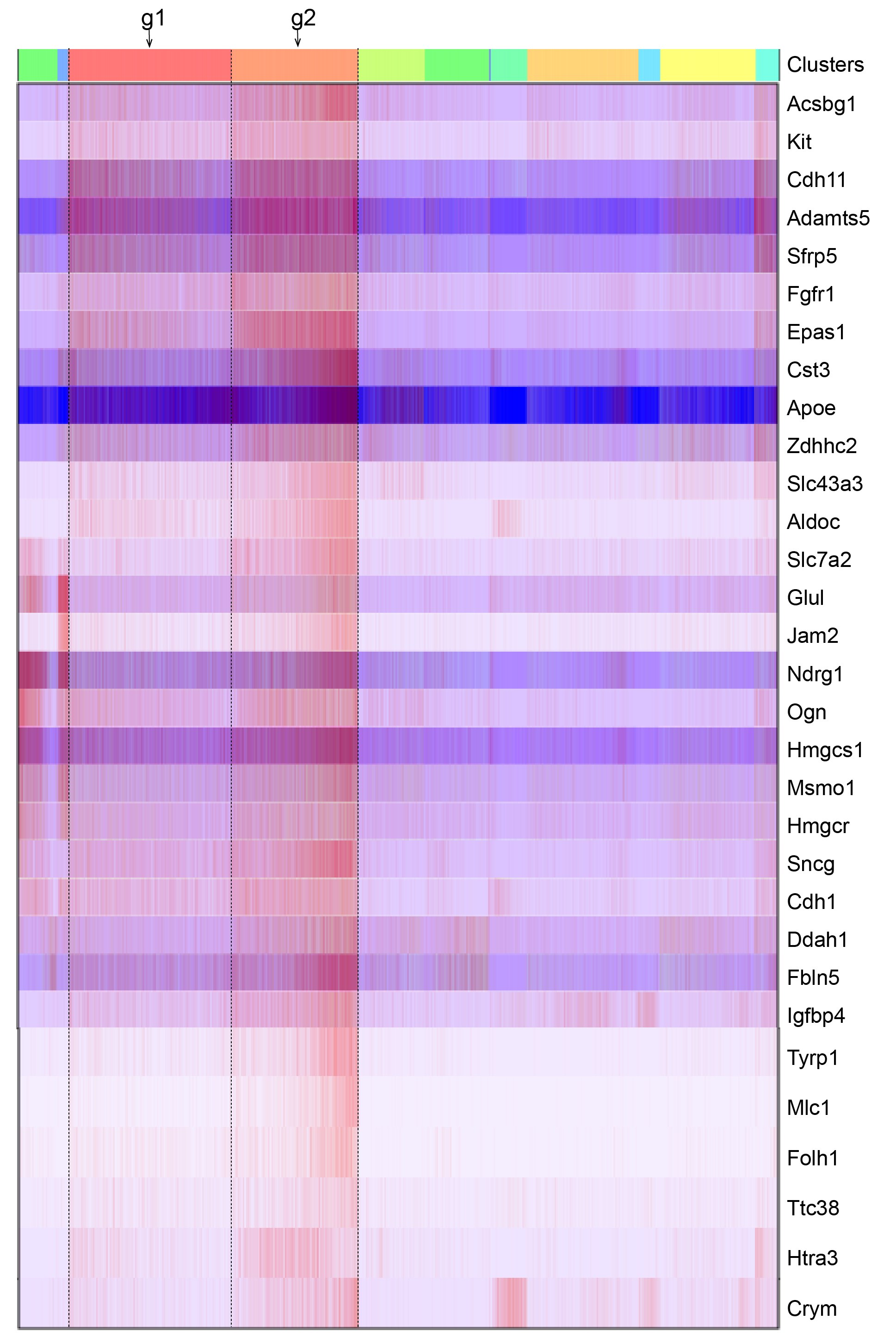

### Figure S9.jpg

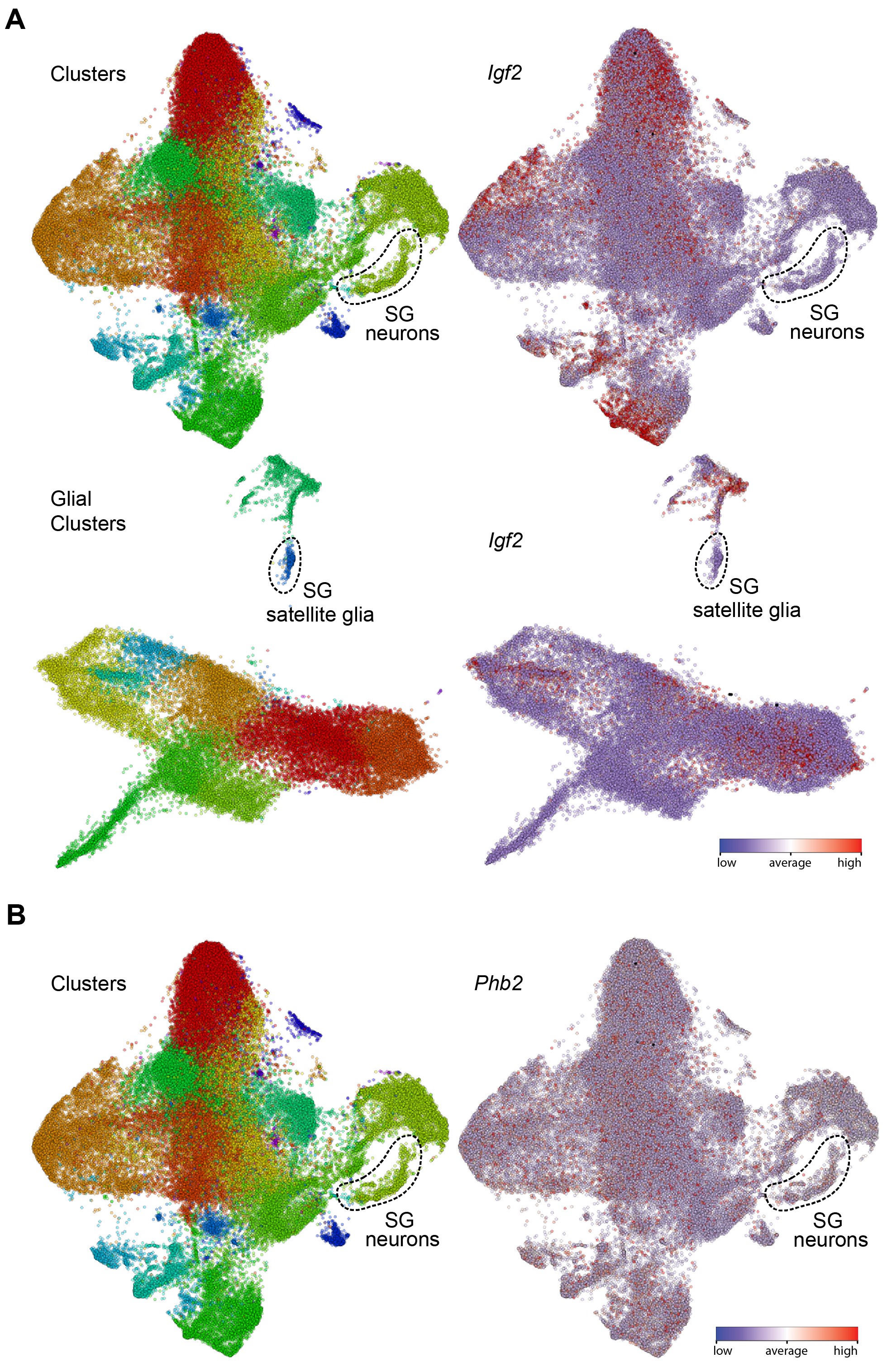

### Figure S10.jpg

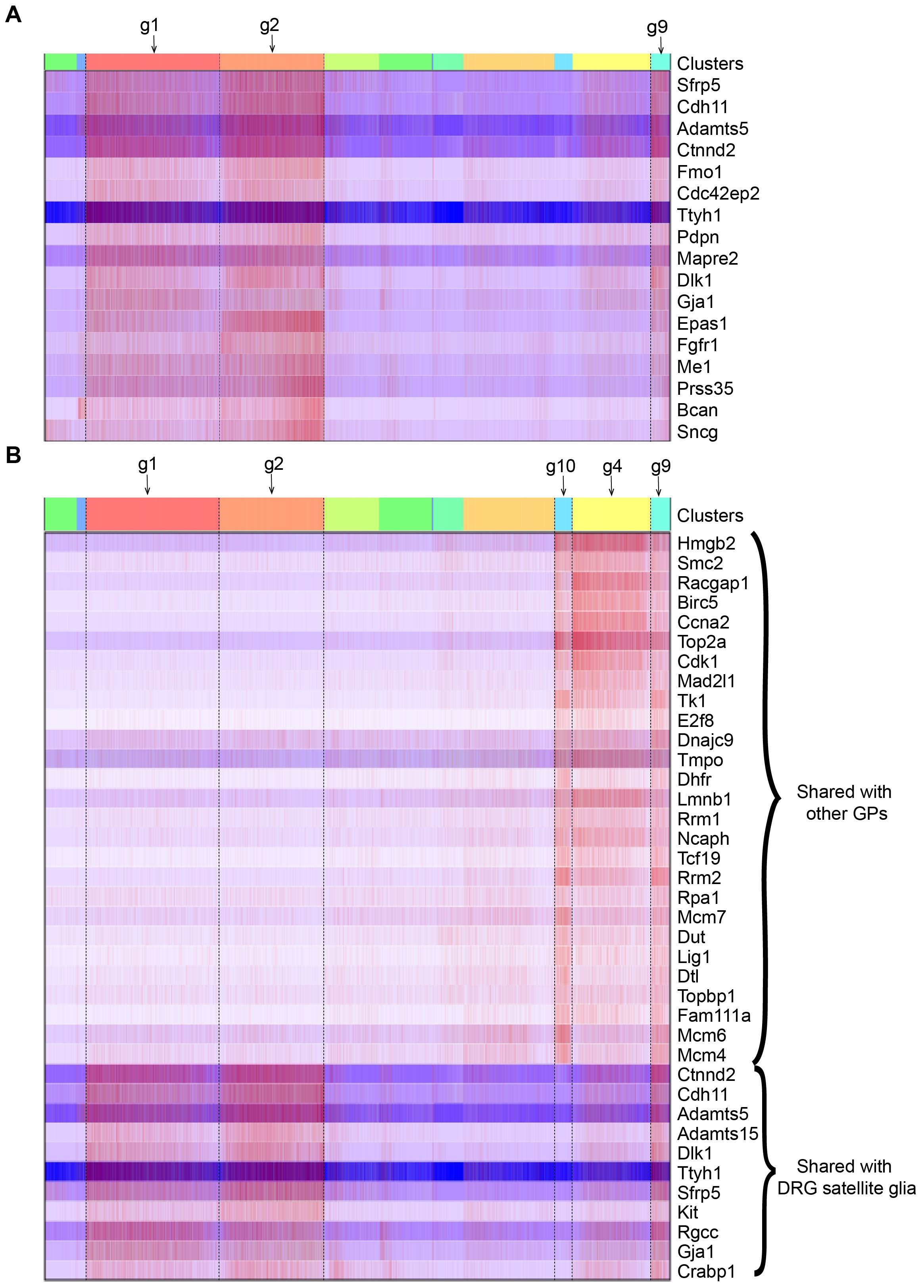

### Figure S11.jpg

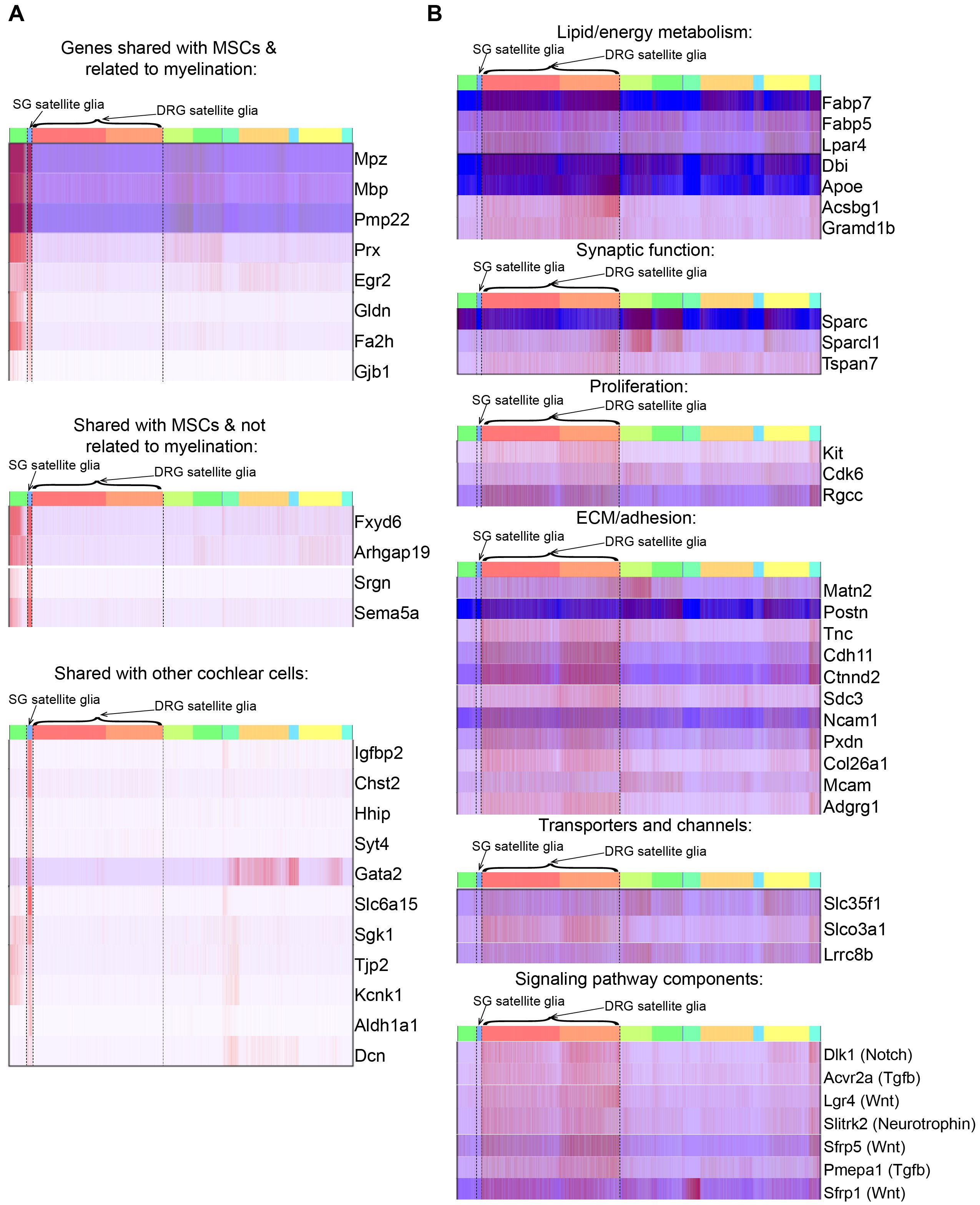
