## Supplemental figure legends for "Diversity of developing peripheral glia revealed by single cell RNA sequencing": Supplemental figure legends.docx

Figure S1: PLP-EGFP is expressed in peripheral glia in both ganglia and supporting cells in cochlea at E14.5. PLP-EGFP labeling glia (A) in lumbar DRG, (B) in cochlea. Schwann cells in the radial bundle projecting towards hair cells are shown with arrows and supporting cells with asterisks. Scale bars: 20um.

Figure S2: The FACS gatings for all the inDrop runs. GFP+ glia are sorted for all. tdtomato+ neurons are not sorted for the P14 run. For E14#1, E14#2, E18#3 and P14 DRG+SN, only the cells collected from the GFP high, not the GFP dim, gating was used for the inDrop collection. For E18#2 cochlea, DN (dominant-negative) was not used for the inDrop collection.

Figure S3: Expression of known markers of non-glial cell types in the dataset. Joint embeddings (UMAP) show expression of markers labeling neurons (Tubb3) (clusters 5 and within 6), SG neurons (Gata3), DRG neurons (Ntrk1), oligodendrocytes (Mobp) (within cluster 6), mesenchymal cells (Pou3f4) (clusters 7 and 11), melanocytes (Mitf) (cluster 13), glia-neuron doublets (Sox10 and Tubb3) (cluster 14), macrophages (Fcgr3) (cluster 15), blood cells (Hba-a1) (cluster 16). Dotted lines surround the indicated cluster.

Figure S4: Expression profile of top genes for GP clusters. Joint embeddings (largeVis) and heatmaps showing expression of *Cdk1*, *Ccna2*, *Birc5* and *Hmgb2*, genes highly expressed in all GP clusters (g4, g9, g10).

Figure S5: GP clusters g9 and g10 predominantly belong to DRG+SN and cochlea, respectively. Joint embeddings (largeVis) highlight cells from (A) g9 in cyan, g9 from cochlea in red and g9 from DRG+SN in blue (B) g10 in light blue, g10 from cochlea in red and g10 from DRG+SN in blue.

Figure S6: Cochlear immature Schwann cells express gene set characteristic of cochlear GPs. Heatmaps show top genes expressed in immature Schwann cell cluster (g3) compared to other glia are also shared with cochlear GPs (g10 and the ones in g4). Curly bracket shows cochlear GPs within the GP cluster g4.

Figure S7: Scn7a is a better marker for NMSCs than common NMSC markers. (A) Joint embeddings (UMAP) and heatmaps of all cells showing L1cam, Ngfr and Scn7a expression. L1cam and Ngfr are expressed in neurons and many glial types, whereas Scn7a is more restricted to NMSC. Neurons and NMSC clusters are surrounded with dotted line. (B) Scn7a levels increase in NMSCs during development. Joint embeddings of one batch from each developmental stage showing Scn7a expression. Arrow points to cells with high Scn7a. (C) RNAscope on P14 sections from *PLP-EGFP* mice shows no Scn7a expression in satellite glia from DRG and SG.

Figure S8: DRG satellite glia cluster g2 shows higher levels of gene expression for satellite glia genes compared to DRG satellite glia cluster g1. Heatmaps of top genes from differential gene expression analysis between clusters g1 & g2, outlined with dotted lines.

Figure S9: Phb2, rather than Igf2, is a candidate molecule expressed in SG neurons to interact with SG satellite glia-expressed Igfbp6. (A) Joint embeddings (UMAP) and (largeVis) show all cells at the top and glial cells at the bottom, respectively. Colors indicate clusters. Igf2 is not expressed in SG neurons nor SG satellite glia compared to other cells. (B) Joint embeddings (UMAP) of all cells. On the left, colors indicate clusters and, on the right, Phb2 expression is seen in SG neurons, indicated with dotted line.

Figure S10: Multiple genes are shared between DRG satellite glia clusters and DRG GP cluster g9. (A) Heatmaps of top genes from differential gene expression analysis between DRG satellite glia (g1 & g2) and other glia show all top genes expressed in g1 & g2 are also expressed in g9. (B) Heatmaps of top genes from differential gene expression analysis between g9 and other glia show a set of genes shared with clusters g1 & g2 in addition to the proliferative gene set shared with GP clusters g4 & g10.

Figure S11: DRG and SG satellite glia show differences in gene expression in multiple categories. (A, B) Heatmaps of top genes (A) expressed higher in cochlear compared to DRG satellite glia, (B) expressed higher in DRG compared to cochlear satellite glia.
