## Supplemental methods for "Diversity of developing peripheral glia revealed by single cell RNA sequencing": STAR METHODS-12_02_20.docx

**KEY RESOURCES TABLE**

| REAGENT or RESOURCE | SOURCE | IDENTIFIER |
| --- | --- | --- |

Antibodies

| Rabbit polyclonal anti mouse Mbp | Abcam | Cat # ab40390 |
| --- | --- | --- |
| Rabbit monoclonal to Ki67 | Vector Laboratories | Cat # VP-RM04 |
| Rabbit polyclonal to GFP | Abcam | Cat # ab6556 |
| Mouse monoclonal anti rat Tuj1 | Biolegend | Cat # 801202 |
| Rat monoclonal anti Mbp | Abcam | Cat# ab7349 |
| chicken anti-Neurofilament | Millipore Sigma | Cat# AB5539 |
| Donkey anti-rabbit, Alexa Fluor 488 | Thermo Fisher Scientific | Cat# A21206 |
| Donkey anti-rabbit, Alexa Fluor 647 | Life Technologies | Cat# A31573 |
| Donkey anti-rabbit, Alexa Fluor 568 | Life Technologies | Cat# A10042 |
| Donkey anti-mouse, Alexa Fluor 647 | Life Technologies | Cat# A31571 |
| Donkey anti-rat DyLight 488 | Novus Biologicals | Cat# NBP1-75383 |
| Donkey anti-chicken, Alexa Fluor 488 | Jackson ImmunoResearch | Cat# 703-545-155 |

Chemicals, Peptides, and Recombinant Proteins

| Collagenase IV | Worthington | Cat# LS004186 |
| --- | --- | --- |
| Papain Dissociation System (Papain, EBSS, ovomucoid inhibitor, DNAse I) | Worthington | Cat# LKLK003150 |
| Trypsin | Worthington | Cat# LS004452 |
| Polystyrene Round-Bottom tube with cell-strainer cap | Falcon | Cat# 352235 |
| Proteinase K | Sigma-Aldrich | Cat# P6556 |
| PFA | Thermo Fisher Scientific | Cat# 50980492 |
| DEPC | Sigma-Aldrich | Cat# D5758 |
| PBS (10X), RNase-free | Life Technologies | Cat# AM9624 |
| Triton X-100 | Millipore | Cat# 648462 |
| ImmEDGE Hydrophobic Barrier Pen | Vector Laboratories | **Cat #** NC9545623 |
| SSC (20X) | ThermoFisher | Cat # 15557-044 |
| Tween-20 | Sigma-Aldrich | Cat# P2287-500ML |
| Prolong Gold antifade reagent | Cell Signaling Technology | Cat # 9071S |
| Donkey serum | Sigma-Aldrich | Cat# D9663 |
| BSA, IgG and protease-free | Jackson ImmunoResearch | Cat# 001-000-162 |
| DAPI | Life Technologies | Cat # D1306 |
| Fluoromount-G | Southern Biotech | Cat# 0100-01 |
| OptiPrep Density Gradient Medium | Sigma-Aldrich | Cat# D1556-250ML |
| HBSS (10X), calcium, magnesium, no phenol red | Life Technologies | Cat# 14065056 |
| HBSS (10X), no calcium, no magnesium, no phenol red | Life Technologies | Cat# 14175103 |
| FBS | Sigma-Aldrich | Cat# F2442 |
| HEPES (1M) | Gibco | Cat# 15630-080 |
| 5X First Strand Buffer | Thermo Fisher Scientific | Cat# 18080-044 |
| Igepal CA-630 | Sigma-Aldrich | Cat# 56741 |
| HFE-7500 oil | Novec | Cat# Novec 7500 |
| MgCl_2_(1M) (nuclease-free) | Thermo Fisher Scientific | Cat# AM9530G |
| DTT | Invitrogen | Cat# 00147 |
| RNaseOUT (40 U/μl) | Thermo Fisher Scientific | Cat# 10777-019 |
| SuperScript III (200 U/μl) | Thermo Fisher Scientific | Cat# 18080-044 |
| Carrier oil | RAN Biotechnologies | Cat# 08-FluoroSurfactant-2wtH-50G |
| Nuclease-free water | Life Technologies | Cat# 10977015 |
| FastDigest Buffer (10X) | Thermo Fisher Scientific | Cat# B64 |
| ExoI (20 U/μl) | Life Technologies | Cat# EN0581 |
| ExoI reaction buffer (10X) | NEB | Cat# B0293S |
| FastDigest HinfI | Thermo Fisher Scientific | Cat# FD0804 |
| Agencourt AMPure XP magnetic beads | Beckman Coulter | Cat# A63881 |
| NEBNext mRNA Second Strand Synthesis Module | NEB | Cat# E6111S |
| HiScribe T7 High Yield RNA Synthesis Kit | NEB | Cat# E2040S |
| RNA Fragmentation Reagents | Ambion/Life Technologies | Cat# AM8740 |
| dNTP (10 mM each) | NEB | Cat# N0447L |
| PrimeScript RT-PCR Kit (including PrimeScript Reverse Transcriptase (200 U/μl) and 5x PrimeScript Buffer) | Takara Clontech | Cat# RR014A |
| Kapa 2X HiFi HotStart PCR mix | Kapa Biosystems | Cat# KK2601 |
| EvaGreen Dye (20X) | Biotium | Cat# 31000-T |
| NextSeq® 500 High Output v2 Kit (75 cycles) | Illumina | Cat# FC-404-2005 |
| BioAnalyzer RNA pico chip and reagents | Agilent | Cat# 5067-1513 |
| Bioanalyzer HS DNA chip and reagents | Agilent | Cat# 50674626 |
| Qubit dsDNA HS Assay Kit | Life Technologies | Cat# Q32851 |
| KAPA Library Quantification Kit | Roche | Cat# 07960140001 |
| EDTA (0.5M) | Thermo Fisher Scientific | Cat# 15575020 |
| Tris-HCl, pH 8.0 (1M) | Thermo Fisher Scientific | Cat# 15568025 |
| Tris-HCl, pH 7.0 (1M) | Thermo Fisher Scientific | Cat# AM9851 |
| Mineral oil | Sigma-Aldrich | Cat# M5310-1L |
| NEG-50 frozen section medium | Richard-Allan Scientific | Cat# 6502 |
| 1H,1H,2H,2H-Perfluorooctanol | Alfa Aesar | Cat# B20156 |
| Ethanol, 200 proof (100%) | Decon Labs | Cat# 22-032-601 |
| Aquapel | Aquapel | Cat# 47100 |
| TSA Plus Cyanine 5 | PerkinElmer | Cat# NEL745E001KT |
| TSA Plus Cyanine 3 | PerkinElmer | Cat# NEL744E001KT |
| Quadrol (N,N,N’,N’-Tetrakis(2-hydroxypropyl)ethylenediamine | Tokyo Chemical Industry | Cat# T0781 |
| Urea | Nacalai Tesque | Cat# 35904-45 |
| Sucrose | Nacalai Tesque | Cat# 30403-55 |
| Triethanolamine (2,2’,2”-Nitrilotriethanol) | Wako | Cat# 145-05605 |

Oligonucleotides

| R1-N6 primer (100 μM) (v3): 5’TCGTCGGCAGCGTCAGATGTGTATAAGAGACAG(N1:25252525) (N1)(N1)(N1)(N1)(N1)3’ | Integrated DNA Technologies | N/A |
| --- | --- | --- |
| R2-PCR primer: 5’CAA GCA GAA GAC GGC ATA CGA GAT GGG TGT CGG GTG CAG3’ | Integrated DNA Technologies | N/A |
| R1-PCR-ix1 primer: 5’AATGATACGGCGACCACCGAGATCTACACCTCTCTATTCGTCGGCAGCGTC3’ | Integrated DNA Technologies | N/A |
| R1-PCR-ix5 primer: 5’AATGATACGGCGACCACCGAGATCTACACAAGGAGTA TCGTCGGCAGCGTC3’ | Integrated DNA Technologies | N/A |
| R1-PCR-ix6 primer:  5’AATGATACGGCGACCACCGAGATCTACACCTAAGCCT TCGTCGGCAGCGTC3’ | Integrated DNA Technologies | N/A |
| R1-PCR-ix8 primer:  5’AATGATACGGCGACCACCGAGATCTACACTCTCTCCG TCGTCGGCAGCGTC3’ | Integrated DNA Technologies | N/A |

Critical Commercial assays

| Hybridization chain reaction (HCR) v3.0 probe hybridization buffer, probe wash buffer and amplification buffer | Molecular Instruments | <https://www.molecularinstruments.com/> |
| --- | --- | --- |
| HCR Probe Set: Mouse Igfbp6 B2 hairpin compatible | Molecular Instruments | PRB540  <https://www.molecularinstruments.com/> |
| HCR Probe Set: Mouse Tubb3 B3 hairpin compatible | Molecular Instruments | PRD 474  <https://www.molecularinstruments.com/> |
| HCR Amplifier B3-h2 Alexa Fluor 546 label | Molecular Instruments | S041721  <https://www.molecularinstruments.com/> |
| HCR Amplifier B2-h2 Alexa Fluor 647 label | Molecular Instruments | S038421  <https://www.molecularinstruments.com/> |
| HCR Amplifier B2-h1 Alexa Fluor 647 label | Molecular Instruments | S041921  <https://www.molecularinstruments.com/> |
| HCR Amplifier B3-h1 Alexa Fluor 546 label | Molecular Instruments | S026521  <https://www.molecularinstruments.com/> |
| RNAscope probe Mm- Ncmap-C2 | Advanced Cell Diagnostics | Cat# 577231-C2 |
| RNAscope probe Mm-Top2a | Advanced Cell Diagnostics | Cat# 491221 |
| RNAscope probe Mm-Scn7a | Advanced Cell Diagnostics | Cat# 548561 |
| RNAscope probe Mm-Epas1-C3 | Advanced Cell Diagnostics | Cat# 314371-C3 |
| RNAscope probe Mm-Igfbp6 | Advanced Cell Diagnostics | Cat# 425721 |
| RNAscope Multiplex Fluorescent Reagent Kit V2 | Advanced Cell Diagnostics | Cat# 323100 |
| Click-iT Edu Cell Proliferation Kit for Imaging, Alexa Fluor 555 dye (containing EdU, Alexa Fluor azide, Click-iT EdU reaction buffer, CuSO_4_, Click-iT Edu buffer additive) | Thermo Fisher Scientific | Cat# C10338 |

Experimental models: Organisms/strains

| Mouse: CD1 (ICR) | Charles River Laboratories | Strain Code: 022 |
| --- | --- | --- |
| Mouse: *PLP-EGFP* | Gabriel Corfas;  (Mallon et al., 2002) | Tg(Plp1-EGFP)10Wmac MGI ID: 5927608 |
| Mouse: *Neurog1^cre^* | Jane Johnston;  (Quinones et al., 2010) | Tg(Neurog1-cre)1Jejo  MGI: 4455179 |
| Mouse: *Neurog1^creERT2^* | Lisa Goodrich;  (Koundakjian et al., 2007) | Tg(Neurog1-cre/ERT2)1Good  MGI: 3777275 |
| Mouse: *Ai14* | The Jackson Laboratory;  (Madisen et al., 2010) | Rosa26-tdTomato  Stock #: 007914 |
| Mouse: *Npy^Cre^* | Qiufu Ma:  (Bourane et al., 2015) | Tg(Npy-cre)RH26Gsat/Mmucd  MMRRC:034810-UCD |

Software and Algorithms

| Conos | <https://github.com/kharchenkolab/conos> |
| --- | --- |

**CONTACT FOR REAGENT AND RESOURCE SHARING**

Further information and requests for resources and reagents should be directed to and will be fulfilled by the Lead Contact, Rosalind Segal.

**EXPERIMENTAL MODEL AND SUBJECT DETAILS**

**Mice**

All animal experiments were approved by the Dana-Farber Cancer Institutional Animal Care and Use Committee and Institutional Animal Care and Use Committee of Harvard Medical School as appropriate and conducted in accordance with the National Institutes of Health guidelines. The following mice lines were used:

1. Single cell RNA-seq experiments: Mice were a result of a cross between the mouse line harboring *PLP-EGFP, Neurog1^cre^*, *Ai14* transgenes or *PLP-EGFP, Neurog1^creERT2^*, *Ai14* and *CD1* line. *Neurog1^cre^* or *Neurog1^creERT2^* along with *Ai14* were used to FACS sort for neurons to spike the glia collection as these mice label sensory neurons in both SG and DRG with *Neurog1^creERT2^*. *Neurog1^creERT2^* without Tamoxifen injection labels sensory neurons sparsely Resulting embryos (E14.5 and E18.5) or pups (P14) were used to dissect lumbar DRGs, SN and cochlea for extracting PLP-EGFP+ glia. Both males and females were used.
2. For fluorescent RNA in situ by HCR or RNAscope methods to validate markers and immunohistochemistry, P14 pups from *PLP-EGFP* line were used.
3. Mice harboring *Npy^Cre^*/+; *Ai14*/+ were used for whole mount immunohistochemistry. Both the *Npy^Cre^* (Bourane et al., 2015) and *Ai14* (Madisen et al., 2010) for this experiment were kindly provided by Dr. Qiufu Ma (Dana-Farber Cancer Institute, Boston, USA).

**METHOD DETAILS**

**Collection of single cells**

Number and genotype of mice dissected for the runs were:

|  | *PLP-EGFP/+* | *PLP-EGFP/+; Neurog1^cre^ /+; Ai14/+* | *Neurog1^cre^ /+* | *PLP-EGFP/+; Neurog1-CreER^T2^/+; Ai14/+* |
| --- | --- | --- | --- | --- |
| 1^st^ E14.5 | 14 | 2 | 1 |  |
| 2^nd^ E14.5 | 17 | 1 |  | 4 |
| 1^st^ E18.5 | 12 | 3 |  |  |
| 2^nd^ E18.5 | 11 | 4 | 2 |  |
| 3^rd^ E18.5 | 17 | 1 | 1 |  |
| 1^st^ & 2^nd^ P14 | 7 | 7 |  |  |

2 cochlea out of the inner ear and 6 lumbar DRGs (3 from each side) and 2 SN (1 from each side) per mouse were dissected. The dissections were done in ice-cold Ca^2+^-free HBSS solution (20mM HEPES in Ca^2+^-free HBSS (Life Technologies, 14175103)). For all dissociations, enzyme mix included collagenase IV, papain and DNase I (from Worthington Papain dissociation system) in HBSS solution with Ca^2+^ (20mM HEPES in HBSS with Ca^2+^ (Life Technologies, 14065056) with only P14 dissociation also included trypsin. The dissociation method was similar for all stages but in order to efficiently get single cells, enzyme concentration and incubation times were changed depending on the stage and the amount of tissue dissected. The aim in increasing enzyme concentration was to minimize the total incubation time as long incubation of cells at 37C can lead to transcriptional changes. As at P14 tissues have increased amounts of myelin and collagen, both enzyme concentration and incubation time had to be increased.

Enzyme concentration and enzyme incubation times for E14, E18, P14:

|  | Enzyme mix | Incubation time |
| --- | --- | --- |
| 1^st^ E14.5 | 20mg/ml Collagenase, 50U/ml papain, 100U/ml DNaseI | 10min |
| 2^nd^ E14.5 | 8mg/ml Collagenase, 50U/ml papain, 100U/ml DNaseI | 25min |
| 1^st^ E18.5 | 8mg/ml Collagenase, 50U/ml papain, 100U/ml DNaseI | 20min |
| 2^nd^ E18.5 | 10mg/ml Collagenase, 50U/ml papain, 100U/ml DNaseI | 15min |
| 3^rd^ E18.5 | 8mg/ml Collagenase, 50U/ml papain, 100U/ml DNaseI | 20min |
| 1^st^ & 2^nd^ P14 | 40mg/ml Collagenase, 100U/ml papain, 750U/ml DNaseI, 2.5mg/ml trypsin | 40min |

Tissues were incubated with the 500ul of the enzyme mix at 37C water bath. After 10min enzyme incubation, trituration with glass pipettes with medium diameter was performed. Then the cells were incubated 5min and triturated with glass with smaller diameter and this is repeated another time with a much smaller pipette to efficiently get single cells. Equal volume of ovomucoid inhibitor in EBSS was then added to stop the reaction. The cells were then spinned with 300rpm for 5-7min, resuspended with 2% BSA in HBSS solution and spinned again to rinse the cells from enzymes and inhibitor. After resuspension with 2% BSA in HBSS solution, the cells were spinned through a tube with cell-strainer cap (Catalog# 352235) and loaded to FACS instrument for sorting. Cell sortings were done at the Dana-Farber Flow Cytometry Jimmy Fund Core at Dana-Farber Cancer Institute. Dead cells were detected by DAPI (1:1000) staining and excluded during FACS. Viable glia (PLP-EGFP+DAPI-) and viable neurons (*Neurog1^cre^*, tdTomato+DAPI-) were sorted. For cochlea glia sample, an additional lower GFP intensity PLP-EGFP+ DAPI- gating is also used to capture more glia. FACS gatings are shown in Figure S2.

At the end of the FACS sorting, the cells were pelleted and resuspended in Optiprep/BSA solution (20% optiprep, 2% BSA in HBSS). An equal percentage of high GFP and low GFP glia from cochlea were combined. For both DRG+SN and cochlea samples, 2.5-5% neurons from the sorting were added to the glial cells to spike the runs as negative control. Cell mixes were loaded at 80,000 cells/ml concentration to the inDrop chip.

**Single cell RNA-seq Cell Capture**

We used inDrops platform to encapsulate and barcode cDNA from single glial cells (Klein et al., 2015; Zilionis et al., 2017). 1^st^ and 2^nd^ E18, 1^st^ E14 and both P14 runs were captured with the Indrop platform we set up following guidelines from Zilionis and colleagues. The materials used for setting up the Indrop platform were listed in Key Resources Table. Hydrogel beads were v3 library format and provided from the Single Cell Core Facility of the ICCB-Longwood Screening Facility at Harvard Medical School. For 2^nd^ E14 and 3^rd^ E18 timepoints, inDrop platform run by the Single Cell Core Facility of the ICCB-Longwood Screening Facility at Harvard Medical School was used. Each sample was divided to aliquots after inDrop collection.

**Library Preparation for next-generation sequencing**

After generating barcoded cDNA from single cells, the droplets were broken in the aliquots, and aliquots were frozen until processed for library preparation. The DRG+SN and cochlea libraries from the same batch were prepared together. The libraries were made following a previously described protocol (Klein et al., 2015; Zilionis et al., 2017), with the following modifications:

- PE2-N6 primer is changed to R1-N6 primer.
- PE1 primer in PE1/PE2 primer mix is changed to R1-PCR-ix primer.
- Multiplexed the libraries with index primers R1-PCR-ix1, -ix5, -ix6 and -ix8 to be able to load two libraries to the sequencer together.
- PE2 primer in PE1/PE2 primer mix is changed to R2-PCR primer.
- The sequencing parameters for next generation sequencing were: Read1: 61 cycles, read2: 14, index 1: 8 cycles, index 2: 8 cycles. We used 5% PhiX as sequencing control. We didn’t use any custom primers.

Libraries were quantified using Qubit and KAPA Library Quantification Kit before next generation sequencing.

For bioinformatic analysis, libraries from two different biological replicates were used for E14, three different biological replicates for E18 experiments and libraries from 2 technical replicates were used for P14.

**Histological Methods**

**mRNA detection by RNAscope**

P14 *PLP-EGFP/+* mice were perfused with 1X PBS followed by 4% PFA. Cochlea, DRG and SN were dissected afterwards and fixed with 4% PFA overnight.

Cochlea tissues were prepared for sectioning as follows: tissues were washed with 1X PBS 2x 10min, decalcified with 120mM EDTA in 1X PBS for 2 days, washed with 1X PBS 2x 10min and then prepared for embedding with 10% sucrose for 30min and 30% sucrose for 3hours and 1:1 mix of 60% sucrose and NEG-50 overnight. Tissues were changed to fresh NEG-50 medium, flash frozen and sectioned at 20um on a cryostat.

DRG and SN tissues were prepared for sectioning as follows: tissues were washed 3x 10min with 1XPBS, then prepared for embedding with 10% to 30% sucrose gradient over 3 days, embedded in NEG-50 medium, and sectioned at 20um on a cryostat.

For RNAscope assays, cochlea, DRG and SN sections were processed together. The manufacturer’s (ACD) protocol for fixed frozen tissues was used with these exceptions:

- The slides were accommodated to room temperature (RT) for 10min as a first step, followed with 10min 1X PBS wash, then post-fix with 4% PFA for 10min and wash with 1X PBS for 2x 5min.
- We did not perform antigen retrieval.
- We incubated tissues with Protease III for 45min at 40C.
- We used TSA Plus fluorophores Cy3 and Cy5 at 1:1750 dilution.
- Immunostain before DAPI addition: tissues were washed with 1X PBS, incubated with block solution (5% donkey serum, 0.1-0.3% tritonX100 in 1X PBS) for 1hr, incubated with rb anti-GFP antibody (1:1000) in block solution overnight at 4C. Next day, tissues were washed with 1X PBS, incubated with donkey anti-rb 488 (1:500) in the same block, RT for 2hrs, washed and mounted following ACD protocol.

**mRNA detection by HCR**

P14 *PLP-EGFP/+* DRG, SN and cochlea sections were prepared as explained for RNAscope. Slides were brought to RT for 10 min, heated at 50C for 15min, fixed with 4% PFA in 1X PBS for 10min at RT, washed with 0.1% Triton X-100 in 1X PBS (PBST) for 2x5min, then treated with 1ug/ml Proteinase K in 1XPBS for 10min at RT, washed with PBST for 2x5min, fixed with 4% PFA in 1X PBS for 10min at RT, washed with PBST for 2x5min, prehybridized with probe hybridization buffer for 10min at 37C, incubated with probes (2ul probe/100ul buffer) at 37C in the humidified chamber overnight. Next day, the hairpins h1 and h2 were snap-cooled (4ul of 3uM stock solutions per slide): heated at 95C on a heat block for 90sec, then cooled to RT in dark. Sections were washed with 100% probe wash buffer for 1min, then with 1.25X SSCT (1XSSCT: 1XSSC, 0.02% Tween 20) in probe wash buffer for 15min at 37C, with 2.5X SSCT in probe wash buffer for 15min at 37C, with 3.75X SSCT in probe wash buffer for 15min at 37C, and with 5X SSCT twice for 15min at 37C. Sections were treated with amplification buffer for 30min at RT, then incubated with hairpin mixtures (3ul/150ul amplification buffer) at RT overnight. Next day, slides were washed with 5X SSCT (first 2x 30min, then 5min) at RT, incubated with block solution (5% donkey serum in 0.1% Triton X-100 in 1X PBS) for 1 hr, then incubated with rabbit anti-GFP ab (1:1000) in block solution at 4C overnight. Next day, the slides were washed with 1X PBS, incubated with secondary ab donkey anti-rb 488 (1:500 in block solution) at RT for 2hrs, washed with 1X PBS, then mounted with Prolong gold antifade.

**Immunohistochemistry on sections**

DRG and cochlea sections from *PLP-EGFP/+* mice at P14 were prepared as above. Sections were brought to RT for 15min, permeabilized with 0.1% Triton X-100 in 1X PBS for 15min, incubated with blocking solution (5% donkey serum, 0.1% Triton X-100 in 1X PBS) for 1hr, incubated with rabbit anti-Mbp (1:200) and mouse anti-Tuj1 (1:500) antibodies in blocking solution at 4C overnight. Next day, slides were washed with 1X PBS, incubated with secondary antibodies donkey anti-rb 568 and donkey anti-mouse 647 (1:500 for both in blocking solution) at RT for 2hrs, washed with 1X PBS, added DAPI, then mounted with Prolong gold antifade.

**Whole mount immunohistochemistry**

2-month-old *Npy^Cre^*/+; *Ai14*/+ mice were used. DRG and SN were dissected and fixed with 4% PFA at 4C overnight, washed 3x 10min with 1X PBS, permeabilized with 0.5% Triton X-100 in 1X PBS for 1hr at RT, incubated with blocking solution (5% donkey serum, 0.5% Triton X-100 in 1X PBS) for 4hr, incubated with rat anti-Mbp (Cat# ab7349) and rabbit anti-Sox2 antibodies (both 1:500) in blocking solution at 4C for 2 days. The tissues were then washed with 1X PBS for a day, then the next day incubated with donkey anti-rat 488 and donkey anti-rabbit 647 in blocking solution (1:500) for 2hrs. Then tissues were washed with 1X PBS and mounted with Fluoromount-G.

For preparation of cochlea, temporal bones were extracted and fixed in 4%PFA overnight. The temporal bones were placed in 120mM EDTA for 2 days with EDTA being changed every day and washed 3x with 1X PBS. Then, the cochlea was dissected out of the temporal bones into three pieces. The cochlea was placed in protease for 30 minutes. Then the cochleae pieces from each mouse were transferred to 30% sucrose for 20 minutes (separate tubes for each mouse). They were then snap frozen on dry ice and quickly thawed by placing the tubes in room temperature water. The tissue was then placed in blocking solution (5% Donkey Serum, 0.3% Triton X-100 in 1X PBS) for 1 hr at RT. The tissues were placed in primary antibody chicken anti-Neurofilament (NF) (Cat# AB5539) 1:1000 in block solution (1% Donkey Serum, 0.3% Triton X-100 in 1X PBS) overnight shaking at 37C. The next day the tissues were washed 3x 15 minutes with 1% TritonX-100 in PBS. The tissues were then incubated with secondary antibody donkey anti-chicken 488 (Cat# 703-545-155) 1:500 in block solution (1% Donkey Serum, 0.3% Triton X-100 in 1X PBS) for 6 hours at 37C, washed 3x 15 minutes with 1% TritonX-100 in PBS. The second wash contained DAPI (10ug/mL). Cochlea were then mounted onto a slide, limbus side down in Fluoromount-G.

Images of DRG and SN were acquired on a Nikon Ni-E C2 confocal with 40X and 60X objectives at 1024x1024 resolution, 16-bit. Stacks were taken from the bottom to the top of the tissues.

Images of the spiral ganglion and radial fibers were acquired on a Leica SP8 confocal with 63X objective at 1024x1024 resolution, 16-bit. Stacks were taken from the bottom to the top of the tissues or single frames were taken at a single plane. Laser power was adjusted so that no channels were being bleached.

**EdU injection and tissue preparation protocol**

Six *PLP-EGFP/+* mice were used at P21. Each mouse was induced with a single dose of 10ug Ethynyl Deoxyuridine (EdU)/gr body weight and was sacrificed one day later. From each mouse, inner ears, lumbar DRGs and SN were dissected after transcardial perfusion in 1XPBS followed by 4% PFA in 1XPBS. Inner ears, DRGs and SN were post-fixed overnight at 4C. After post-fix, DRGs and SN sections were prepared as described for mRNA detection protocols above. After post-fix, inner ears were washed 2x with 1xPBS for 10 min, then decalcified in 120mM EDTA for 2 days, changing the solution at the beginning and end of each day. Then, inner ears were embedded in NEG-50 frozen section medium and sectioned at 16-20um on a cryostat.

**EdU detection and Ki67 immunohistochemistry on sections**

For EdU detection, Click-iT Edu Cell Proliferation Kit for Imaging, Alexa Fluor 555 dye is used. Stock solutions for EdU reaction were prepared following the kit manual. Slides were brought to RT for 15min to dry, rinsed with 1XPBS for 10min, post-fixed with 4%PFA for 10min, rinsed with 1X PBS for 10min, washed with 3% BSA in PBS for 10min, permeabilized with 0.5% Triton X-100 in PBS for 20min and washed 2x with 3% BSA in PBS. incubated with Click-iT reaction cocktail, a mix of reaction buffer, CuSO4, Alexa Fluor azide, reaction buffer additive from the kit, was prepared following kit manual. Slides were incubated with Click-iT reaction cocktail for 30min in dark, washed with 3% BSA in PBS, then incubated with block solution (5% donkey serum in 0.1% Triton X-100 in 1X PBS) for 2hrs, incubated with rb anti-Ki67 antibody (1:500 in block solution) at 4C overnight. Next day, slides were washed with 1X PBS, incubated with secondary ab donkey anti-rb 647 (1:500 in block solution) at RT for 2hrs, washed with 1X PBS, then mounted with Prolong gold antifade.

**CUBIC clearing of cochlear tissue sections**

The *PLP-EGFP*/*+* E14.5 cochlea tissue sections are cleared by CUBIC method modified from (Susaki et al., 2015). Animals are perfused with cold, freshly prepared 4% PFA followed by fixing the whole head overnight with 4% PFA. After the brain is removed, it is washed 3x with 1X PBS then sectioned on vibratome (300-1000um). The sections were washed with ½ Reagent-1 (10% Triton, 5% NNN-tetrakis (2-HP) ethylenediamine, 10% Urea, 25mM NaCl in ddH20) for 30 minutes, then with Reagent-1 for 4 hours, rinsed up to 5x with 1X PBS and left in 1X PBS overnight. For clearing, sections were washed with ½ Reagent-2 (25% urea, 50% sucrose, 10% triethanolamine in ddH20) in 1X PBS for 30 minutes, then washed with Reagent-2 for 3-4 hours.

Then, the sections were pre-incubated in mineral oil for 10-30 minutes and mounted on slide in hybridization chamber.

**QUANTIFICATION AND STATISTICAL ANALYSIS**

**Bioinformatic analysis**

The demultiplexing and barcode analysis was performed using InDrops pipeline (https://github.com/indrops/indrops), using Genome Reference Consortium Mouse Build 38 (GRCm38) to estimate the molecular count matrix for the measured cells. The count matrices were integrated using Conos (<https://github.com/kharchenkolab/conos>), using PCA rotation space (ncomp=30) with angular distance measure, estimated on top 2000 overdispersed genes (n.odgenes=2000) and the neighborhood radius *k*=15. Potential doublets were evaluated using scrublet algorithm (Wolock et al., 2019), a cluster of doublet cells (p>0.25) was removed, and integration re-ran. UMAP and largeVis embeddings were used for the full (Figure 1C) and glial-only (Figure 1F) integrations, respectively. Joint clusters were generated using Leiden clustering algorithm. Differential expression was carried out using Wilcoxon rank sum test, as implemented in Conos. In Figures, the gene expression was visualized by taking log-expression vector: $x_{g}=log(c_{g}*\frac{{10}^{3}}{S}+1)$, where $c_{g}$ is the number of molecules of a gene *g* in a given cell, and *S* is the total number of molecules in the cell. To visualize relative expression, $x_{g}$ was trimmed at 10% using Winsorization procedure, centered by the mean, and normalizing by the $max(\left| x_{g} \right|)$.

**Edu and Ki67 immunofluorescence quantification**

The images were taken on a Nikon Ni-E C2 confocal with 40X objective and images were blinded afterwards. PLP-EGFP+ Edu+ and PLP-EGFP+ Ki67+ cells were counted from each image by a second individual blinded to genotype. The volume is calculated from the area and thickness of the confocal images using Fiji (Image J). Area is calculated by using the drawing tool and the whole tissue are is traced.

**Scn7a immunofluorescence quantification**

The images were taken on a Nikon Ni-E C2 confocal with 60X objective. PLP-EGFP+ Scn7a+ cells were counted from each RB and SN image that were stained by RNAscope for Scn7a probe. The Scn7a+ glia counts are normalized to total DAPI count. Total DAPI count was performed using Fiji (Image J) software: maximum intensity projections were created from confocal stacks, thresholded with Li method, segmented with watershed plugin and 100um-inf range was used to select for DAPI+ nuclei from background speckles.

**DATA AND SOFTWARE AVAILABILITY**
